# Integrative small and long RNA-omics analysis of human healing and non-healing wounds discovers cooperating microRNAs as therapeutic targets

**DOI:** 10.1101/2021.12.17.473103

**Authors:** Zhuang Liu, Letian Zhang, Maria A. Toma, Dongqing Li, Xiaowei Bian, Irena Pastar, Marjana Tomic-Canic, Pehr Sommar, Ning Xu Landén

## Abstract

MicroRNAs (miR) are important posttranscriptional regulators and exhibit a high potential to be utilized in diagnosis and therapy. However, our insufficient knowledge of the miR-mediated gene regulation in human skin wound healing severely hinders the identification of clinically relevant miRs. Here, we performed paired small RNA and long RNA sequencing in human tissue samples, including matched skin and acute wounds collected at each healing stage and chronic non-healing venous ulcers (VU). With integrative small and long RNA-omics analysis, we developed a compendium (https://www.xulandenlab.com/humanwounds-mirna-mrna), which will be an open, comprehensive resource to broadly aid wound healing research. With this first clinical, wound-centric resource of miRs and mRNAs, we identified 17 pathologically relevant miRs that exhibited abnormal VU expression and displayed their targets enriched explicitly in the VU gene signature. Intermeshing regulatory networks controlled by these miRs revealed their high cooperativity in contributing to chronic wound pathology characterized by persistent inflammation and proliferative phase initiation failure. Furthermore, we demonstrated that miR-34a, miR-424, and miR-516, upregulated in VU, cooperatively suppressed keratinocyte growth while promoting inflammatory response. Collectively, our study opens the possibility of developing innovative wound treatment that targets pathologically relevant cooperating miRs to attain higher therapeutic efficacy and specificity.

## Introduction

Wound healing is a fundamental biological process comprising three sequential and overlapping phases, i.e., inflammation, proliferation, and remodeling(Reinke & Sorg, 2012). This delicate repair process is often disrupted in chronic venous insufficiency patients, resulting in venous ulcers (VU) characterized by persistent inflammation and proliferative phase initiation failure(Eming et al., 2014). VU is the most common chronic non-healing wound type, comprising 45–60% of all lower extremity ulcerations(Vivas et al., 2016). VU exhibits a marked impact on health-related life quality and represents a significant financial burden both to the patients and the society with an annual health care cost of overall $14.9 billion in the USA(Hoversten et al., 2020). A deeper understanding of the underlying gene expression regulatory mechanisms during physiological and pathological wound repair is essential for developing more effective wound treatments(Stone et al., 2017).

MicroRNAs (miR) represent a group of short (∼22 nt) non-coding ribonucleic acids, incorporating into the RNA-induced silencing complex and binding to the 3’ untranslated region of their target mRNAs, resulting in mRNA destabilization and translational repression(Stavast & Erkeland, 2019). Given that an individual miR can target dozens to hundreds of genes, miRs have been identified as regulators of complex gene networks(Stavast & Erkeland, 2019). MiR-mediated regulation is reportedly crucial in multiple fundamental biological processes including skin wound repair(Herter & Xu Landen, 2017; Meng et al., 2018). Importantly, manipulating miRs critical for the disease pathogenesis could offer a prominent therapeutic effect, supported by viral infection- and cancer-targeting miR therapeutics clinical trials(Rupaimoole & Slack, 2017). Therefore, miR-based therapeutics for hard-to-heal wounds represent a promising approach(Herter & Xu Landen, 2017; Luan et al., 2018; Meng et al., 2018; Nie et al., 2020; Pastar et al., 2021; Sen & Roy, 2012).

However, our insufficient knowledge of the miR-mediated gene regulation in human wounds severely hinders the identification of clinically relevant miRs and their potential therapeutic use. While most previous wound healing-related miR studies rely on *in vitro* or animal models, only a few have approached miR profiles in human wound tissues or primary cells from patients, including tissues and fibroblasts of diabetic foot ulcers(Liang et al., 2016; Ramirez et al., 2018), burn wound dermis(Liang et al., 2012), and acute wounds at the inflammatory phase(Li et al., 2015). Despite sharing several fundamental features, the human skin structure and repair processes are different from those of the commonly used animal models (e.g., rodents)(Elliot et al., 2018). Moreover, animal models cannot fully simulate the human disease complexity, and the findings are difficult to extrapolate to humans(Darwin & Tomic-Canic, 2018; Pastar et al., 2018). Thereby, a rigorous and in-depth characterization of miR-mediated gene regulatory networks in human healing and non-healing wounds is timely needed.

In this study, we performed paired small and mRNA expression profiling in the human skin, acute wounds during the inflammatory and proliferative phases, and VU, unraveling time-resolved changes of the whole transcriptome throughout the wound healing process and the unique gene expression signature of a common chronic wound type. The integrative miR and mRNA omics analysis provides a network view of miR-mediated gene regulation in human wounds *in vivo* and demonstrates the functional involvement of miRs in human skin wound repair at the system level. Importantly, we identified miRs highly relevant to VU pathology, based not only on their aberrant expression but also their targetome enriched in the VU-related gene expression signature. Apart from confirming the *in silico* findings, the experimental miR expression, targetome, and function validation uncovered that VU- dysregulated miRs could act cooperatively contributing to the stalled wound healing characterized by failed transition from inflammatory-to-proliferative phase, which opens up new possibility for the development of more precise and innovative wound treatment targeting pathologically-relevant cooperating miRs to achieve higher therapeutic efficacy and specificity. Additionally, based on this comprehensive analysis of human wound tissues, we built a browsable resource web portal (https://www.xulandenlab.com/humanwounds-mirna-mrna), which is the first wound healing-focused miR resource for facilitating the exploration of miR’s clinical application and for aiding in the elucidation of posttranscriptional regulatory underpinnings of tissue repair.

## Materials and Methods

### Human wound samples

Human wound biopsies were obtained from 10 healthy donors and 12 patients with chronic venous ulcer (VU) at the Karolinska University Hospital Solna (Stockholm, Sweden). Donor demographics are presented in **Table 1**. Patients with VUs, which persisted for more than four months despite conventional therapy, were enrolled in this study (**Table 2**). Tissue samples were collected from the lower extremity at the nonhealing edges of the ulcers by using a four-mm biopsy punch (**Figure 1a**). Healthy donors above 60 years old without skin diseases, diabetes, unstable heart disease, infections, bleeding disorder, immune suppression, and any ongoing medical treatments were recruited (**Table 3**).

**Figure 1.**
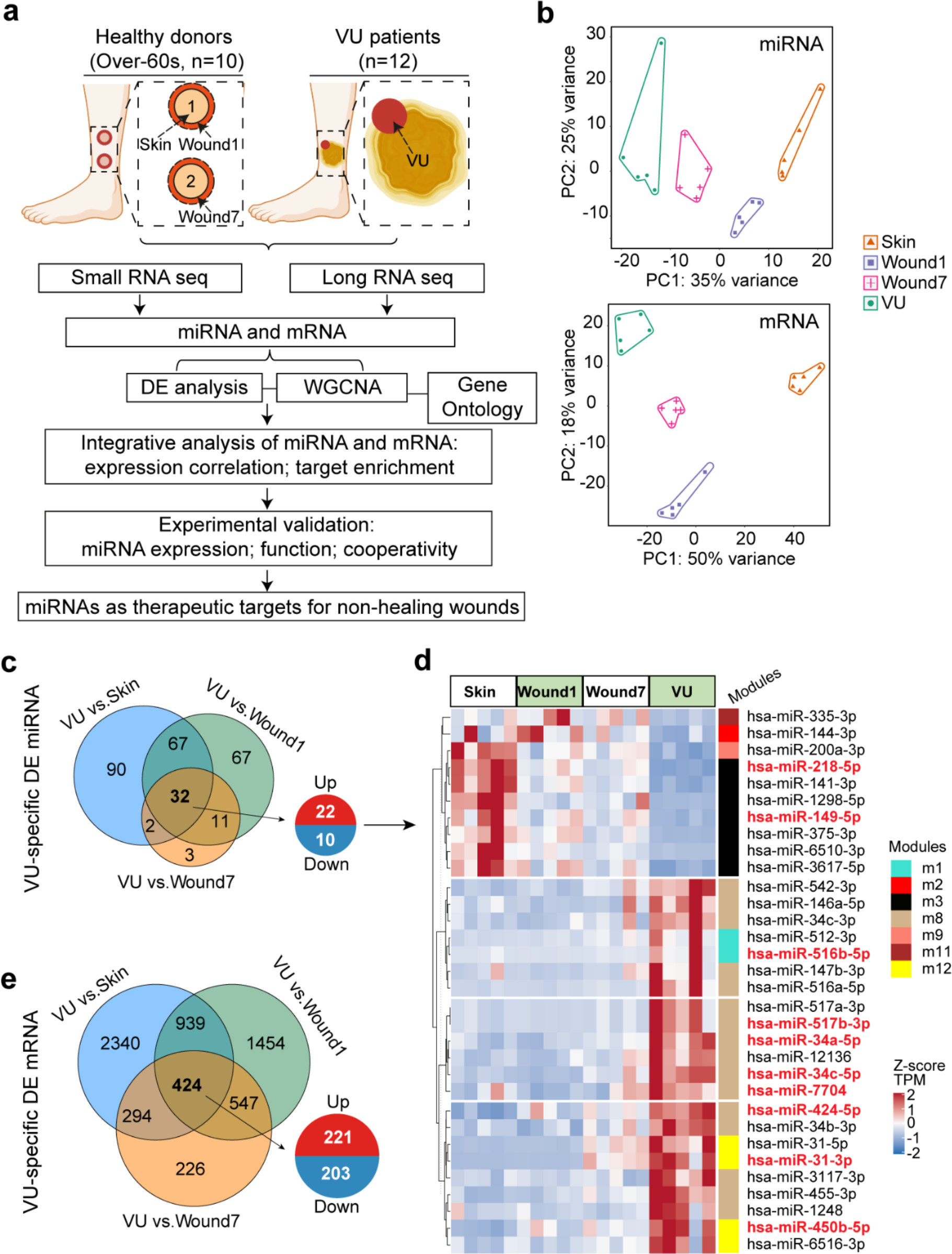
Paired profiling of miRNA and mRNA expression in human wounds. a Schematic of analysis in this study. **b** PCA plots based on miRNA (upper panel) and mRNA (lower panel) expression profiles. Each dot indicates an individual sample. The numbers of differentially expressed (DE) miRNAs **c** and mRNAs **e** in VU (n=5) compared to the Skin, Wound1, and Wound7 from 5 healthy donors are shown in Venn diagrams. FDR < 0.05, fold change ≥ 2 for miRNAs and ≥ 1.5 for mRNAs. **d** The heatmap depicts the 32 miRNAs specifically dysregulated in the VU with scaled expression values (Z-scores). WGCNA modules of each miRNA belongs to are marked with color bars. The miRNAs with experimentally validated expression changes are highlighted in red.

**Table 1.** Characteristics of the healthy donors and the VU patients.

| **Characteristics** | **Healthy donors** | **Patients with VU** |
| --- | --- | --- |
| Study population (n) | 10 | 12 |
| Age, years (mean ± s.d.) | 65.3 ± 3.2 | 75.8 ± 12.0 |
| Ethnicity | White | White |
| Gender (male: female) | 2:8 | 5:7 |
| Biopsy location | Lower leg | Lower leg |
| Wound duration | Acute (1 or 7 days after injury) | 3.7 ± 5.3 years |

Abbreviations: s.d., standard deviation; VU, venous ulcer.

**Table 2.** Characteristics of the patients with venous ulcer.

| Patient | Sex | Age (years) | Ethnicity | Wound size (cm) | Wound duration (years) | Wound location | Experiment |
| --- | --- | --- | --- | --- | --- | --- | --- |
| 1 | M | 86 | Caucasian | 6x5 | 4 | Lower leg | RNA-seq & qRT-PCR |
| 2 | F | 68 | Caucasian | 15x15 | 2 | Lower leg | RNA-seq & qRT-PCR |
| 3 | M | 70 | Caucasian | 3x0.5 | 0.3 | Lower leg | RNA-seq & qRT-PCR |
| 4 | F | 78 | Caucasian | 15x12 | 1.5 | Lower leg | RNA-seq & qRT-PCR |
| 5 | F | 87 | Caucasian | 3x2+8x4 | 2.5 | Lower leg | RNA-seq & qRT-PCR |
| 6 | F | 73 | Caucasian | 3x4 | 3.5 | Lower leg | qRT-PCR |
| 7 | M | 99 | Caucasian | 20x10 | 0.5 | Lower leg | qRT-PCR |
| 8 | F | 71 | Caucasian | 20x20 | 20 | Lower leg | qRT-PCR |
| 9 | F | 77 | Caucasian | 2.5x3+15x15 | 4.5 | Lower leg | qRT-PCR |
| 10 | M | 51 | Caucasian | 2x1.5 | 3 | Lower leg | qRT-PCR |
| 11 | M | 69 | Caucasian | 12x15 | 1 | Lower leg | qRT-PCR |
| 12 | F | 81 | Caucasian | 7x2.5 | 1 | Lower leg | qRT-PCR |

**Table 3.** Characteristics of the healthy donors.

| Donor | Sex | Age<br>(years) | Ethnicity | Wound location | Experiment |
| --- | --- | --- | --- | --- | --- |
| 1 | F | 66 | Caucasian | Lower leg | RNA-seq |
| 2 | M | 69 | Caucasian | Lower leg | RNA-seq |
| 3 | F | 67 | Caucasian | Lower leg | RNA-seq |
| 4 | M | 69 | Caucasian | Lower leg | RNA-seq & qRT-PCR |
| 5 | F | 64 | Caucasian | Lower leg | RNA-seq & qRT-PCR |
| 6 | F | 60 | Caucasian | Lower leg | qRT-PCR |
| 7 | F | 66 | Caucasian | Lower leg | qRT-PCR |
| 8 | F | 60 | Caucasian | Lower leg | qRT-PCR |
| 9 | F | 67 | Caucasian | Lower leg | qRT-PCR |
| 10 | F | 65 | Caucasian | Lower leg | qRT-PCR |

1084 Abbreviations: M, male; F, female.

Two full-thickness excisional wounds (4 mm in diameter) were created at the lower extremity on each donor, and the excised skin was saved as intact skin control (Skin). The wound-edges were excised with a six mm-biopsy punch at day one (Wound1) and day seven (Wound7) after wounding (**Figure 1a**). Written informed consent was obtained from all the donors to collect and use the tissue samples. The study was approved by the Stockholm Regional Ethics Committee and conducted according to the Declaration of Helsinki’s principles.

### RNA extraction, library preparation, and sequencing

#### RNA extraction

Snap frozen tissue samples were homogenized with the TissueLyser LT (Qiagen), and total RNA was isolated using the miRNeasy Mini kit (Qiagen). RNA quality and quantity were determined by using Agilent 2100 Bioanalyzer (Agilent Technologies) and Nanodrop 1000 (Thermo Fisher Scientific Inc.), respectively.

#### Small RNA library preparation and sequencing

The small RNA sequencing libraries were constructed using 3 μg total RNA per sample and NEB Next® Multiplex Small RNA Library Prep Set for Illumina® (NEB) following the manufacturer’s recommendation. Briefly, total RNA was first ligated to adaptors at the 3’ end by NEB 3’ SR adaptor and 5’ end by T4 RNA ligase followed by reverse transcription into cDNA using M-MuLV Reverse Transcriptase. PCR amplification of cDNA was performed using SR primers for Illumina and index primers. The PCR products were purified, and DNA fragments spanning from 140 to 160bp were recovered and quantified by DNA High Sensitivity Chips on the Agilent Bioanalyzer. The libraries were sequenced on an Illumina Hiseq 2500 platform (Illumina, Inc.) using single-end 50bp reads, and all samples were run side by side.

#### mRNA library preparation and sequencing

The long RNA sequencing libraries were constructed with a total amount of 2 μg RNA per sample. First, the ribosomal RNA was depleted by Epicentre Ribo-zero® rRNA Removal Kit (Epicentre). Second, strand-specific total-transcriptome RNA sequencing libraries were prepared by incorporating dUTPs in the second-strand synthesis step with NEB Next® Ultra^TM^ Directional RNA Library Prep Kit for Illumina® (NEB). Finally, the libraries were sequenced on an Illumina Hiseq 4000 platform, and 150 bp paired-end reads were generated for the following analysis.

### Analysis of miRNA-sequencing data

#### Quality control, mapping, and quantification

Quality of raw data was assessed using FastQC v0.11.8 (http://www.bioinformatics.babraham.ac.uk/projects/fastqc/). We used mapper.pl module in the miRDeep2 v0.1.3 package(Friedlander et al., 2012; Mackowiak, 2011) to filter low-quality reads and remove sequencing adaptors and redundancies. Trimmed reads with lengths greater than 18 nucleotides were mapped to GENCODE human reference genome (hg38) by the software Bowtie v1.2.2(Langmead et al., 2009). The miRDeep2.pl module was then performed with default parameters to identify known miRNAs, which were compared to miRNAs in miRBase release 22.1(Kozomara et al., 2018). Counts of reads mapped to each known mature miRNAs were acquired from the quantifier.pl module output without allowing mismatch. miRNAs with read counts less than five in more than half of twenty samples were discarded since these miRNAs are unlikely to give stable and robust results. Raw counts of 562 miRNAs were normalized for sequencing depth using TPM methods (transcript per million = mapped read count/total reads * 10e6)(Zhou et al., 2010) and prepared for further analysis.

#### Differential expression (DE) analysis

The DESeq2 workflow(Love et al., 2014) was carried out to fit raw counts to the negative binomial (NB) generalized linear model and to calculate the statistical significance of each miRNA in each comparison. In particular, the paired model was employed when comparing samples from the same donor. *P-value*s obtained from the Wald test were corrected using Benjamini-Hochberg (BH) multiple test to estimate the false discovery rate (FDR). The differentially expressed miRNAs were defined as FDR < 0.05 and |log2(fold change)| ≥ 1.

#### Principal component analysis (PCA)

To explore the similarity of each sample, PCA was performed by using a DESeq2 built-in function *plotPCA* on the transformed data, in which the variances and size factors were stabilized and corrected. PCA and heatmaps were plotted by using *ggplot2*(Hadley, 2016) and *ComplexHeatmap*(Gu et al., 2016) packages in RStudio (https://rstudio.com/).

#### Weighted gene co-expression network analysis (WGCNA)

The normalized expression of 562 miRNAs were used as input to the WGCNA R package(Langfelder & Horvath, 2008). First, we calculated the strength of pairwise correlations between miRNAs using the ‘biweight’ mid-correlation method. The function *pickSoftThreshold* was then employed to compute the optimized soft-thresholding power based on connectivity, which led to an approximately scale-free network topology(Zhang & Horvath, 2005). Second, a signed weighted co-expression network was constructed with a power of 18 using the one-step *blockwiseModules* algorithm (**Figure 2 - figure supplement 2a**). Network modules were filtered according to parameters: *minModuleSize*= 10 and *mergeCutHeight* = 0.25.

**Figure 2.**
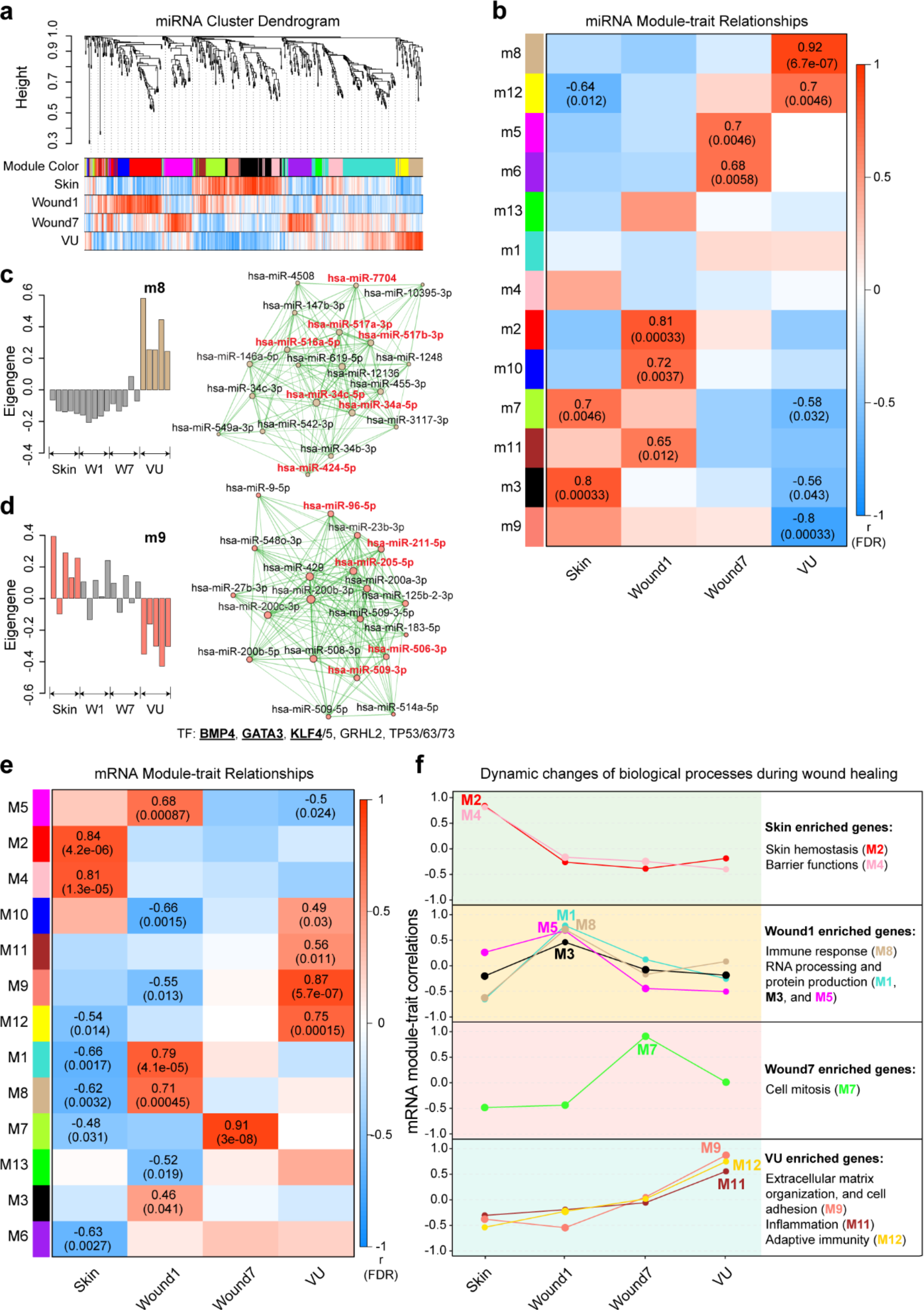
Weighted gene co-expression network analysis (WGCNA) of miRNAs and mRNAs in wound healing. a. Cluster dendrogram shows miRNA co-expression modules: each branch corresponds to a module, and each leaf indicates a single miRNA. Color bars below show the module assignment (the 1^st^ row) and Pearson correlation coefficients between miRNA expression and the sample groups (the 2^nd^ to the 5^th^ row: red and blue lines represent positive and negative correlations, respectively). **b** Heatmap shows Pearson correlations between miRNA module eigengenes (ME) and the sample groups. The correlation coefficients and the adjusted *P-values* (FDR) are shown where the FDRs are less than 0.05. For the VU-associated modules m8 **c** and m9 **d**, bar plots (left) depict the ME values across the 20 samples analyzed by RNA-seq, and network plots (right) show the top 20 miRNAs with the highest kME values in each module. Node size and edge thickness are proportional to the kME values and the weighted correlations between two connected miRNAs, respectively. The miRs with their targetome enriched with VU-mRNA signature (see Figure 4b) are highlighted in red. Transcription factors (TFs) with their targets enriched in the m9 module (Fisher’s exact test: FDR < 0.05) are listed below the network, and TFs differentially expressed in VU are underlined. **e** Heatmap shows Pearson correlations between mRNA MEs and the sample groups. **f** The gene expression pattern of each module across all the sample groups is depicted with line charts. Gene ontology analysis of biological processes enriched in each module is shown at the right.

The expression profile of each module was represented by the module eigengene (ME), referred to as the first principal component of all miRNAs in each module. Pearson correlations (values from -1 to 1) and the corresponding *P-value*s between MEs and traits were computed. *P-value*s were further adjusted to FDR across all the modules using the BH method. Modules significantly associated with each trait were selected by FDR < 0.05 and absolute correlation coefficients > 0.4. The module membership (also known as kME) of each miRNA was calculated by the correlations between miRNA expression and ME. MiRNAs with the highest kME values were defined as intra-modular hub miRNAs, and networks of hub miRNAs in significant modules were visualized using the Cytoscape v3.7.2 software(Shannon et al., 2003).

To check the robustness of module definition, we carried out module preservation analysis and calculated the standardized Z-scores for each module by permutating 200 times using the same 20 samples as reference and test datasets. Modular preservation is strong if Z-summary > 10, weak to moderate if 2 < Z-summary < 10, no evidence of preservation if Z-summary ≤ 2(Langfelder et al., 2011).

#### Transcription factor (TF) enrichment analysis

We leveraged a curated database about TF-miRNA regulations, TransmiR v2.0(Tong et al., 2019), to identify the TFs regulating miRNA expression in each module. Fisher’s exact tests were employed to evaluate the enrichment of each TF in the significant modules, and FDRs were adjusted to the total number of TFs (Odds ratio > 1 and FDR < 0.05). Correlations of gene expression between TFs and miRNA modules (represented by MEs) were further filtered to identify putative TF-mediated miRNA gene expression patterns (Pearson correlation: *P-value* < 0.05, coefficient > 0). The 562 miRNAs abundantly expressed in our samples were treated as the background dataset.

### Analysis of mRNA sequencing data

Raw reads of mRNA sequencing were first trimmed for adaptors and low-quality bases using Trimmomatic v0.36 software(Bolger et al., 2014). Clean reads were aligned to the human reference genome (GRCh38.p12), coupled with the comprehensive gene annotation file (GENCODEv31) using STAR v2.7.1a(Dobin et al., 2013). Gene expression was then quantified by counting unique mapped fragments to exons by using the feature count function from the Subread package(Liao et al., 2013). Raw counts for each gene were normalized to fragments per kilobase of a transcript, per million mapped reads (FPKM)-like values. Only mRNAs with FPKM ≥ 1 in at least ten samples were kept for the rest analysis. We used the same pipeline described above for mRNA DE and PCA analysis. The differentially expressed mRNAs were defined as FDR < 0.05 and |log2(fold change)| ≥ 0.58. WGCNA was carried out for 12,069 mRNAs with the optimal threshold power of 12 according to a fit to the scale-free topology of the co-expression network (**Figure 2 - figure supplement 4a**). Thirteen mRNA modules were identified with the settings: *maxBlockSize* = 20000, *minModuleSize* = 100 and *mergeCutHeight* = 0.25. Furthermore, mRNA module-enriched TF analysis was performed with a manually curated TF-target regulatory relationship database, TRRUST v2(Han et al., 2018), using Fisher’s exact tests. TFs with FDR < 0.05 and odds ratio > 1 and the expression significantly correlated with respective mRNA modules (Pearson correlation *P-value* < 0.05) were identified.

### Gene ontology (GO) analysis

We carried out GO analysis for mRNAs by using the WebGestalt tool (http://www.webgestalt.org/) (Liao et al., 2019), which applied a hypergeometric test in target and reference gene sets. GO terms of non-redundant biological process (BP) with gene number less than 10 and adjust *P- value* (FDR) > 0.05 were filtered out.

### Integrative analysis of miRNA and mRNA expression changes in wound healing

#### Expression correlation between mRNA and miRNA modules

An integrative analysis was carried out by relating the first principal component (PC1) of miRNA expression, calculated using the *moduleEigengenes* function of WGCNA package(Langfelder & Horvath, 2008), to the PC1 of mRNA expression in each module. The miRNA-mRNA module pairs with a Pearson correlation coefficient < -0.5 and a *P-value* < 0.05 were selected for the following enrichment analysis.

#### Prediction of miRNA targets

We predicted both conserved and non-conserved target sites for all the 562 miRNAs by using the *get_multimir* function from R package multimiR(Ru et al., 2014) (http://multimir.org/) based on the latest TargetScan v7.2 database(Agarwal et al., 2015; Lewis et al., 2005). All predicted miRNA targets were sorted by a primary score calculated for target site strength, and the top 25% with summed context++ score ≤ -0.15 were defined as the strongest miRNA targets. Targets that were not detected by the long RNA-seq were removed.

#### Gene set enrichment analysis of miRNA targets in mRNA modules

We evaluated the degree of enrichment of miRNA modules’ targets in mRNA modules. For this, we focused on the VU-specific DE miRNA, i.e., the 22 up- and 10 down-regulated miRNAs in VU compared to both the skin and acute wounds (FDR < 0.05 and |log2(fold change)| ≥ 1), as well as the VU-associated modules’ hub miRNAs, which kME values were greater than the median kME in respective modules (i.e., 14 miRNAs in m8, 9 miRNAs in m12, 20 miRNAs in m7, 29 miRNAs in m3, and 13 miRNAs in m9). Among these miRNAs’ strongest targets, we selected the ones that were hit by ≥ 2 miRNAs from m8, m9, m12 modules or miRNAs downregulated in VU; ≥ 3 miRNAs from m7 module or miRNAs upregulated in VU; ≥ 4 miRNAs from m3 module, to capture putative module-driving targets. We performed gene set enrichment analyses for these miRs’ targets in VU-specific DE mRNAs (FDR < 0.05 and fold change ≥ 1.5) and VU-associated mRNA modules by using the R function *fisher.test()* based on the two-side Fisher’s exact test(Wu et al., 2016). Furthermore, we performed enrichment analysis to identify individual miRNA with their strongest targets significantly enriched in VU-specific DE mRNAs or VU-associated mRNA modules (Fisher’s exact test: odds ratio > 1, *P-value* < 0.05).

### Experimental validation of miRs’ expression, targetome, and functions

#### Quantitative RT-PCR

To detect miRNA, RNA from human skin and wounds was reverse transcribed using TaqMan® Advanced miRNA cDNA Synthesis Kit (Thermo Fisher Scientific). Individual miRNA expression was then quantified using TaqMan® Advanced miRNA Assays (Thermo Fisher Scientific) and normalized with miR-361-5p and miR-423-5p due to their relatively constant expression between human skin and wounds. To detect mRNA, we performed reverse transcription using the RevertAid First Strand cDNA Synthesis Kit (ThermoFisher Scientific). Gene expression was examined by SYBR Green expression assays (ThermoFisher Scientific) and normalized with housekeeping gene B2M and GAPDH. The primer sequences for B2M are forward primer (5’-AAGTGGGATCGAGACATGTAAG-3’) and reverse primer (5’-GGAGACAGCACTCAAAGTAGAA-3’); GAPDH forward primer (5’- GGTGTGAACCATGAGAAGTATGA-3’) and reverse primer (5’-GAGTCCTTCCACGATACCAAAG-3’).

#### Primary cell culture and transfection

Adult human epidermal keratinocytes were cultured in EpiLife serum-free keratinocyte growth medium supplemented with Human Keratinocyte Growth Supplement (HKGS) and 100 units/mL Penicillin and 100 μg/mL Streptomycin (Thermo Fisher Scientific). Adult human dermal fibroblasts were cultured in Medium 106 supplemented with Low Serum Growth Supplement (LSGS) and 100 units/mL Penicillin and 100 μg/mL Streptomycin (Thermo Fisher Scientific). Cells were incubated at 37°C in 5% CO2, and media was routinely changed every 2–3 days. Third passage keratinocytes at 50%-60% confluence were transfected with 20 nM miRNA mimics (Horizon) or negative control using Lipofectamine^TM^ 3000 (Thermo Fisher Scientific).

#### Microarray analysis

Transcriptome profiling of keratinocytes and fibroblasts transfected with 20 nM miRNA mimics or control mimics for 24 hours (in triplicates) was performed using Affymetrix Genechip system at the Microarray Core facility of Karolinska Institute. Normalized expression data (log2 transformed value) were exported from Transcriptome Analysis Console (TAC) software and analyzed by the *limma* R package(Ritchie et al., 2015). In brief, expression data were first fitted to a linear model for each probe. Then, the empirical Bayes method was applied to compute the estimated coefficients of gene-wise variability and standard errors for comparisons of experimental and control groups. Genes with FC > 1.2 and *P*-value < 0.05 between the miRNA mimics- and the control mimics-transfected cells were considered to be significantly changed. Gene set enrichment analysis, including biological process (BP), Kyoto Encyclopedia of Genes and Genomes (KEGG) pathway, and hallmark from Molecular Signatures Database (MSigDB) (http://www.gsea-msigdb.org/)(Subramanian et al., 2005), was performed with a ranked fold change list of all the genes by using the *fgsea* R package(Korotkevich et al., 2021) and visualized by using the *ggplot2*(Hadley, 2016) and *circlize*(Gu et al., 2014) packages.

#### Immunofluorescence staining

Cells transfected with a combination of miRNA and/or control mimics (50 nM in total) were fixed in 4% paraformaldehyde (PFA) for 15 minutes. Cells were incubated with the Ki- 67 antibody (Cell Signaling Technology) overnight at 4°C. The next day, cells were incubated with the secondary antibody conjugated with Alexa 488 for 40 minutes at room temperature. Cells were mounted with the ProLong™ Diamond Antifade Mountant with 4’,6-Diamidino-2-Phenylindole (DAPI) (ThermoFisher Scientific). Ki-67 signals were visualized with Nikon microscopy, and positive cells were counted using ImageJ software (National Institutes of Health).

#### Cell proliferation assay

Cells were seeded in 12-well plates with a density of 20,000 cells/well. The plates were placed in IncuCyte live-cell imaging and analysis platform (Essen Bioscience) after cells attaching to the plates. Plates were imaged every two hours, and pictures were processed and analyzed using IncuCyte ZOOM 2018A software (Essen Bioscience).

#### Statistical analysis

Sample size of each experiment is indicated in the figure legend. Data analysis was performed by using R and Graphpad Prism 7 software. Comparison between two groups was performed with Mann-Whitney U tests (unpaired, non-parametric), Wilcoxon signed-rank test (paired, non- parametric), or two-tailed Student’s t-test (parametric). The cell growth assay was analyzed by using two-way ANOVA. *P*-value < 0.05 is considered to be statistically significant.

## Results

### miRNA and mRNA paired expression profiling in human wounds

To better understand tissue repair in humans, we collected wound-edge tissues from human acute wounds and chronic non-healing VUs (**Figure 1a** and **Table 1–3**). We created 4mm full thickness punch wounds at the lower legs of healthy volunteers aged beyond 60 years to match the advanced age of VU patients and anatomical location of the highest VUs occurrence(Vivas et al., 2016). Tissue was collected at baseline (Skin), and at day one and day seven post-wounding (Wound1 and Wound7) to capture the inflammatory and proliferative phases of wound healing, respectively. In total, 20 samples divided into four groups, i.e., Skin, Wound1, Wound7, and VU, were analyzed by Illumina small RNA sequencing (sRNA-seq) and ribosomal RNA-depleted long RNA sequencing (RNA-seq). After stringent raw sequencing data quality control (**Table S1** and **S2**), we detected 562 mature miRs and 12,069 mRNAs in our samples. Our principal component analysis showed that either the miR or the mRNA expression profiles clearly separated these four sample groups (**Figure 1b**). Next, we performed pairwise comparisons to identify the differentially expressed genes (DEG) during wound repair. We compared the VUs with both the skin and acute wounds and unraveled a VU-specific gene signature, including aberrant increase of 22 miRs and 221 mRNAs and decrease of 10 miRs and 203 mRNAs (DE analysis FDR < 0.05, fold change ≥ 2 for miRs and ≥ 1.5 for mRNAs, **Figure 1c–e** and **Additional file 1**). The full DEG list can be browsed on our resource website (https://www.xulandenlab.com/humanwounds-mirna-mrna) with more or less rigorous cut-offs. With this unique resource, we dissected further the miR-mediated posttranscriptional regulatory underpinnings of wound repair.

### Dynamically changed miR expression during wound repair

We leveraged weighted gene co-expression network analysis (WGCNA) for classifying miRs according to their co-expression patterns in the 20 sRNA-seq-analyzed samples to link the miR expression changes with wound healing progression or non-healing status at a system level(Langfelder & Horvath, 2008). We identified 13 distinct modules with a robustness confirmed by the module preservation analysis (**Figure 2 - figure supplement 1a**), ten of them significantly correlating (Pearson’s correlation, FDR < 0.05) with at least one of the four phenotypic traits, i.e., Skin, Wound1, Wound7, and VU (**Fig. 2a** and **b, Additional file 2**). The WGCNA revealed that module (m)2, m10, and m11 miRs were upregulated at the inflammatory phase (Wound1), while m5 and m6 miRs peaked at the proliferative phase (Wound7). In VU, we identified three downregulated (m3, m7, and m9) and two upregulated (m8 and m12) miR modules. We highlighted the 198 “driver” miRs (i.e, the top 20 miRNAs with the highest kME values in each module and kME > 0.5) of the ten significant modules in the co-expression networks (**Figure. 2c, d** and **Figure 2 - figure supplement 2b–i**) and they could also be browsed on our resource web portal (https://www.xulandenlab.com/humanwounds-mirna-mrna). Notably, we identified 84% of them as DEGs, suggesting a high consistence between the WGCNA and DE analysis (**Additional file 3**).

We hypothesized that the co-expression of various miRs could be due to their transcription driven by common transcription factors (TF). To test this idea, we leveraged TransmiR v2.0(Tong et al., 2019), a database including literature-curated and ChIP-seq-derived TF-miR regulation data, to identify the enriched TFs in each miR module [Fisher’s extract test: odds ratio (OR) > 1, *FDR* < 0.05, **Additional file 4**]. Interestingly, the BMP4, KLF4, KLF5, GATA3, GRHL2, and TP53 families exhibited not only their binding sites enriched in the m9 miR genes but their expression also significantly correlated with the m9 miRs (Pearson’s correlation coefficient = 0.53–0.82, *P*-value of *p* = 7.05e-06–0.014) (**Figure 2d, Figure 2> - figure supplement 3a** and **Additional file 4**). Notably, the BMP4, GATA3, and KLF4 expressions were significantly reduced in VU compared to the skin and acute wounds. This result could explain the deficiency of their regulated miRs in VU and also suggest a link between these TFs and chronic wound pathology (**Figure 2 - figure supplement 3b**).

### mRNA co-expression networks underpinning wound repair

miRs exert functions through the posttranscriptional regulation of their target mRNAs. Therefore, describing the mRNA expression context would be required for understanding the role of miRs in wound repair(Agarwal et al., 2015). We thus performed WGCNA in the paired long RNA-seq data and identified 13 mRNA co-expression modules (**Figure. 2e**, **Figure 2 - figure supplement 1b**, **Figure 2 - figure supplement 4a–c**, and **Additional file 5**). The GO analysis of the mRNA modules largely confirmed the previous knowledge of wound biology, such as skin hemostasis (M2) and barrier function (M4)- related gene downregulation in the wounds, the upregulation of the genes involved in the immune response (M8), RNA processing, and protein production (M1, M3, and M5) in the inflammatory phase, and the prominent cell mitosis-related gene expression (M7) in the proliferative phase of wound repair (**Figure. 2f** and **Figure 2 - figure supplement 5a**). These results further supported the robustness and reproducibility of our profiling data. Moreover, this unique dataset allows the identification of the key TFs driving these biological processes. For example, we identified NFKB1 and RELA, well-known for their immune functions(Liu et al., 2017), as the most enriched upstream regulators for the M1 mRNAs, while E2F1, a TF promoting cell growth(Ertosun et al., 2016), surfaced as a master regulator TF in M7 (**Additional file 6**).

Importantly, our study unraveled a VU molecular signature: downregulated expression of RNA and protein production- (M1, M3, and M5) as well as cell mitosis-related (M7) genes, and upregulated expression of genes involved in extracellular matrix organization and cell adhesion (M9). These results were in line with the dermal tissue fibrosis observed in patients with chronic venous insufficiency(Blumberg et al., 2012; Pappas et al., 1999; Stone et al., 2020). Moreover, we found an immune gene signature clearly distinguishing the chronic inflammation in VUs (M11 and M12 enriched with adaptive immunity-related mRNAs) from the self-limiting immune response in acute wounds (M8 enriched with neutrophil activation- and phagocytosis-related mRNAs) (**Figure 2f** and **Figure 2 - figure supplement 5b**). Overall, we generated a gene expression map of human healing and non-healing wounds, setting a stepping stone for the in-depth understanding of the VU pathological mechanisms. After having established this map, we decided to dissect how miRs contribute to these pathological changes.

### Integrative analysis of miR and mRNA expression changes in wound healing

Among the multiple gene expression regulatory mechanisms, we aimed to evaluate how miRs could contribute to the protein-coding gene expression in human wound repair. We thus performed a correlation analysis for the miRs and mRNAs that were differentially expressed in VU compared to the skin and acute wounds, using the first principal component (PC1) of their expression in each sample. We found significantly negative correlations (Pearson’s correlation, P-values: 1.36e-12–1.27e-04) between the PC1 of the DE miRs and the DE mRNAs predicted as miR targets, indicating negative regulation of VU-mRNA signature by the aberrantly expressed miRs in VU (**Figure 3a–c**).

**Figure 3.**
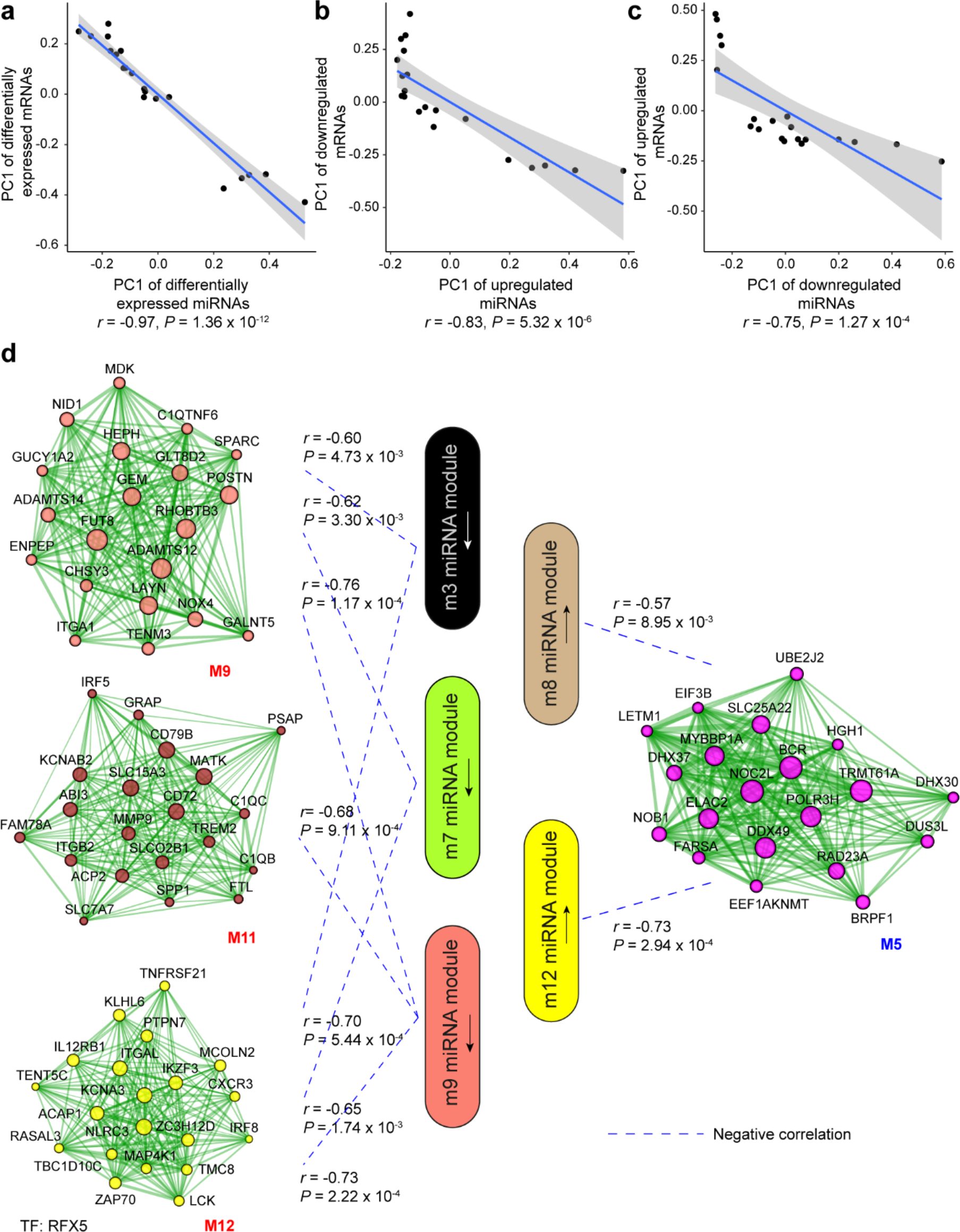
Correlation analysis between miRNA and mRNA expression changes in VU. a-c. Correlations between the first principal component (PC1) of VU-associated differentially expressed (DE) miRNAs, and the PC1 of VU DE mRNAs predicted as miRNA targets. **d** PC1 correlations between the hub miRNAs and their predicted targets in the VU-associated miRNA and mRNA modules. Pearson correlation coefficients (*r*) and *P* values are shown. The mRNA networks are plotted with the top 20 most connected module genes. Transcription factors (TFs) with targets enriched in the VU-associated modules are listed below the networks.

Furthermore, we dissected the potential regulatory relationship between the VU-associated miR and mRNA modules. We identified significantly negative correlations between the downregulated miR (m3, m7, and m9) and the upregulated mRNA (M9, M11, and M12) modules, as well as between the upregulated miR (m8 and m12) and the downregulated mRNA (M5) modules in VU (**Figure 3d**). Among these miR-mRNA module pairs, we found that the predicted targets of the downregulated m9 miRs were significantly enriched (Fisher’s extract test: OR > 1, *P*-value < 0.05) for the upregulated M9 mRNAs, whereas the targets of the upregulated miRs and m8 miRs were enriched for the downregulated mRNAs and M5 mRNAs (**Figure 4a** and **Additional file 7**). These results demonstrated that miRs significantly contribute to the aberrant mRNA expression in VU at a global level.

**Figure 4.**
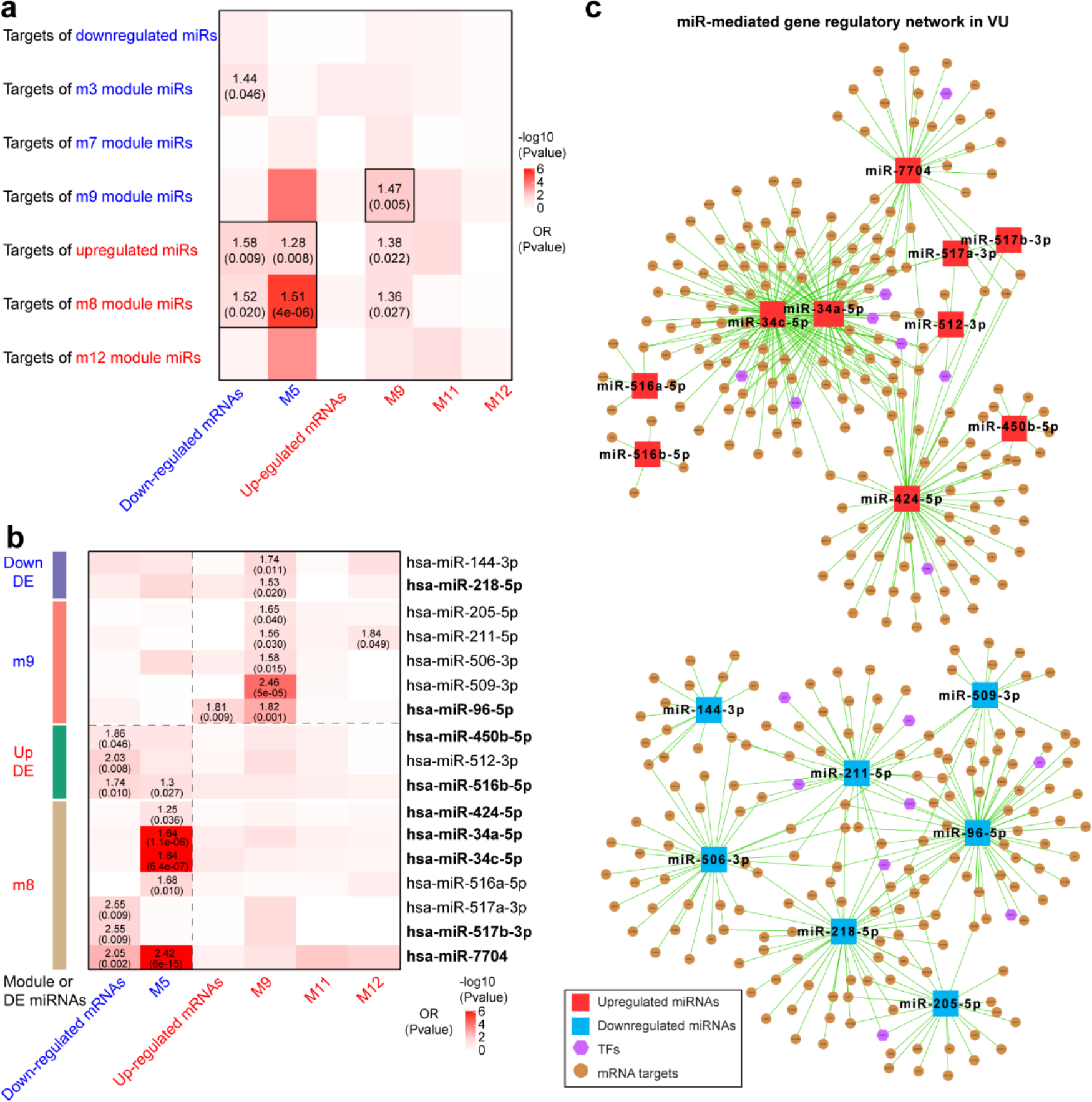
Integrative analysis of miRNA and mRNA expression changes in VU. Heatmaps show the enrichment for VU-affected mRNAs and mRNA modules in the top targets of **a** VU-associated DE miRNAs and miR modules and **b** the individual candidate miRNAs. Odds ratio (OR) and *P* values are indicated when OR > 1 and *P-value* < 0.05 (Fisher’s exact test). The miRNAs with experimentally validated expression changes are highlighted in bold. **c** miR-mediated gene regulatory networks in VU are constructed with the miRNAs in Figure 4b, the mRNAs predicted as the strongest targets and with anti-correlated expression patterns (Pearson correlation, *P-value* < 0.05 and r < 0) with these miRNAs in human wounds, and the TFs regulating these miRNAs’ expression from the TransmiR v2 database. An enlarged version of these networks can be found in Figure 4 – figure supplement 1 and 2.

Based on the above-identified miR-mRNA module pairs, we next searched for individual candidate miRs with their targets enriched for the VU mRNA signature. We observed that the targets of two VU-associated downregulated miRs (miR-144-3p and miR-218-5p) and five m9 miRs (miR-205-5p, miR- 211-5p, miR-506-3p, miR-509-3p, and miR-96-5p) were enriched for the upregulated M9 mRNAs, whereas the targets of three VU-associated upregulated miRs (miR-450-5p, miR-512-3p, and miR- 516b-5p) and seven m8 miRs (miR-424-5p, miR-34a-5p, miR-34c-5p, miR-516a-5p, miR-517a-3p, miR- 517b-3p, and miR-7704) were enriched for M5 mRNAs and downregulated mRNAs (**Figure 4b** and **Additional file 8**). These miR targetomes were enriched for the mRNAs associated with VU pathology. Therefore, these miR candidates are of importance for understanding the pathological mechanisms hindering wound healing. Moreover, we compiled miR-mediated gene expression regulation networks centered with these highly pathologically relevant miRs (**Figure 4c**, **Figure 4 – figure supplement 1**, **2** and **Additional file 8**). These networks also include the mRNAs predicted as the strongest targets and with anti-correlated expression patterns with these miRs in human wounds *in vivo*, as well as the TFs reported to regulate these miR expressions from the TransmiR v2 database(Tong et al., 2019). Taken together, our study identifies a list of highly pathological relevant miRs and their targetomes in human VU.

### Experimental validation of miR expressions and targets in human skin wounds

We selected nine shortlisted DE miRs (**Figure 1d** and **4b**), including three downregulated (miR-149-5p, miR-218-5p, and miR-96-5p) and six upregulated (miR-7704, miR-424-5p, miR-31-3p, miR-450-5p, miR-516b-5p, and miR-517b-3p) miRs in VU, and validated their expression by qRT-PCR in a cohort with seven healthy donors and twelve VU patients, matched in terms of age and the anatomical wound locations (**Table 2** and **3**). We confirmed their expression patterns in RNA-seq, supporting the robustness and reproducibility of our profiling data (**Figure 5a–i**, **Figure 5 – figure supplement 1 and Additional file 9**).

**Figure 5.**
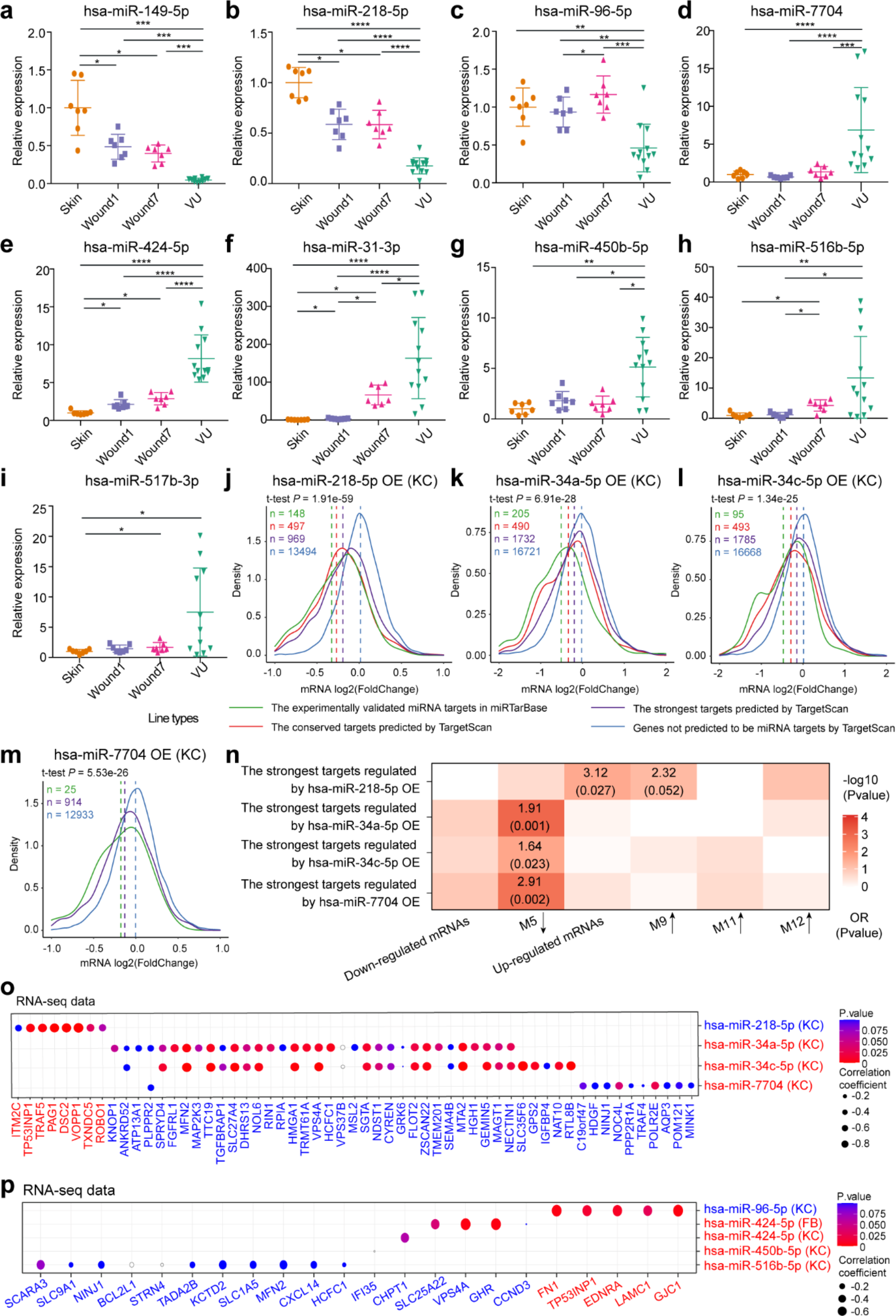
Experimental validation of miRNAs’ expression and targets in human skin wounds. a-i qRT-PCR analysis of VU-associated DE miRNAs in the skin, day 1 and day 7 acute wounds from 7 healthy donors and venous ulcers (VU) from 12 patients. Wilcoxon signed-rank test was used for the comparison between Skin, Wound1, and Wound7; Mann-Whitney U test was used for comparing VU with the skin and acute wounds. \**P* < 0.05, \*\**P* < 0.01, \*\*\**P* < 0.001, \*\*\*\**P* < 0.0001. **j-m** Microarray analysis was performed in keratinocytes (KC) with miR-218-5p (**J**), miR-34a-5p **k**, miR-34c-5p **l**, or miR- 7704 **m** overexpression (OE), and density plots of mRNAs log2(fold change) are shown. Wilcoxon t- tests were performed to compare the TargetScan predicted strongest targets (purple) with the non- targets (blue) for each of these miRNAs. The conserved and experimentally validated targets are marked with red and green colors, respectively. Dotted lines depict the average log2(fold change) values for each mRNA group. **n** A heatmap shows the enrichment for VU-affected mRNAs and mRNA modules in the targets of miR-218-5p, miR-34a-5p, miR-34c-5p, or miR-7704 validated by the microarray analysis. Odds ratio (OR) and *P* values are shown when OR > 1 and *P-value* < 0.05 (Fisher’s exact test). **o, p** For each of the miRNAs (miR-218-5p, miR-34a-5p, miR-34c-5p, miR-7704, miR-96, miR-424, miR-450b, and miR-516b) and its targets validated by the microarray analysis of KC or fibroblasts (FB), Pearson correlation analysis was performed between their expression values in human skin and wound samples shown by RNA-seq. Grey circles indicate correlation coefficients > 0.

Furthermore, we experimentally validated the targets of eight miRs surfaced in our analysis (**Figure 4b**), including the miRs downregulated (miR-218-5p and miR-96-5p) and upregulated (miR-424-5p, miR- 450-5p, miR-516b-5p, miR-34a-5p, miR-34c-5p, and miR-7704) in VU. We performed genome-wide microarray analysis in human primary keratinocytes or fibroblasts overexpressing each of these miRs. Furthermore, we re-analyzed our published microarray dataset on keratinocytes with miR-34a-5p or miR-34c-5p overexpression (GSE117506)(Pachera et al., 2020). For all these eight miRs, we observed that their strongest targets predicted by TargetScan were significantly downregulated compared to the non-targeting mRNAs (Wilcoxon t-test P-values: 1.34e-25∼1.91e-59). The differences were more significant when we divided the strongest targets into conserved and experimentally validated subtypes, confirming the bioinformatics prediction robustness of the miR targets applied in this study (**Figure 5j– m** and **Figure 5 – figure supplement 2a - j**).

Notably, we observed significant enrichment (Fisher’s exact test, OR > 1, *P*-value *<* 0.05) of the experimentally validated miR-218-5p, miR-34a-5p, miR-34c-5p, and miR-7704 targets for the VU gene signature (**Figure 5n** and **Additional file 9**). We have previously shown that miR-34a and miR-34c inhibit keratinocyte proliferation and migration, while promoting apoptosis and inflammatory response, resulting in delayed wound repair in a mouse model(Pachera et al., 2020). We validated robustness of the bioinformatics approach applied in this study by miR-34a/c re-identification. Here, we discovered that both the predicted (**Figure 4b**) and validated (**Figure 5n**) miR-34a/c targets were enriched for the downregulated M5 module mRNAs in VU. Notably, our microarray analysis confirmed that miR-34a/c reduced the expression of 39 hub genes in the M5 module [log2(fold change) ≤ -0.58, *P*-value < 0.05, Fig. S9j], and 26 of them exhibited negative correlation (Pearson’s *r* = −0.83 – −0.45, *P*-value < 0.05) with miR-34a/c expression levels in the human skin and wound samples (**Figure 5o** and **Additional file 9**), suggesting that they were miR-34a/c targets *in vivo*. MiR-218-5p was downregulated in the VU compared to the acute wounds and the skin (**Figure 5b** and **Figure 5 – figure supplement 1**). Its predicted (**Figure 4b**) and validated (**Figure 5n**) targets were both enriched for the upregulated or M9 module mRNAs in VU. Among the ten *in vitro* validated targets, eight negatively correlated (Pearson’s *r* = −0.82 ∼ −0.46, *P*-value of *p* < 0.05) with miR-218-5p expression in the human skin and wounds (**Figure 5o**, **Figure 5 – figure supplement 2j**, and **Additional file 9**). Interestingly, this study also identified miR-7704, a human-specific miR, with significantly increased expression in VU (**Figure 5d** and **Figure 5 – figure supplement 1**). Similar to miR-34a/c, the predicted (**Figure 4b**) and validated (**Figure 5n**) miR-7704 targets were highly enriched for the M5 module mRNAs downregulated in VU.

For miR-96-5p, miR-424-5p, miR-450-5p, and miR-516b-5p, although their predicted targets were significantly enriched (Fisher’s exact test: OR > 1, *P*-value < 0.05) for VU-associated mRNAs (**Figure 4b**), we did not find similar enrichment for their experimentally validated targets. Nevertheless, these miRs regulated some VU-associated hub genes *in vitro* (**Figure 5 – figure supplement 2k**) and also exhibited an anti-correlated expression pattern with their targets *in vivo* (**Figure 5p**), e.g., miR-96- 5p from the downregulated m9 module targets the M9 mRNAs upregulated in VU, including TP53INP1, LAMC1, EDNRA, GJC1, and FN1; while miR-424-5p from the upregulated m8 miR module targets the M5 mRNAs downregulated in VU, including SLC25A22, VPS4A, and GHR (**Additional file 9**).

In summary, we experimentally validated the expression and targets of the miRs identified by the RNA-seq data bioinformatics analysis, confirming the robustness and reproducibility of this dataset and highlighting its value as a reference for studying the physiological and pathological roles of miRs in human skin wound healing.

### Cooperativity of VU pathology-relevant miRs

From the miR-mediated gene expression regulation networks underpinning VU pathology (**Figure 4 – figure supplement 1, 2,** and **Additional file 8**), we caught a glimpse of presumable miR cooperativity through targeting the same mRNAs, i.e., co-targeting among miRs, which reportedly imposing stronger and more complex repression patterns on target mRNA expression(Cherone et al., 2019). For the miRs with unrelated seed sequences, we found that miR-34a/c and miR-424-5p or miR-7704 shared eight– ten targets, and these miRs were co-expressed in the m8 module. We showed that among the downregulated miRs in VU, miR-96-5p and miR-218-5p shared eight targets.

In addition, we performed functional annotations for the genes regulated by the VU-associated miRs identified in the microarray analysis (**Figure 6a** and **Additional file 10**). Both miR-218-5p and miR-96- 5p promoted ribosome biogenesis and non-coding (nc) RNA processing, while miR-218-5p also suppressed keratinization. miR-34a/c-5p enhanced innate immune response, while reducing mitosis. Similarly, miR-424-5p and miR-516b-5p increased the cellular defense response, while inhibiting cell proliferation. In addition, we showed that miR-450-5p upregulated genes related to the ncRNA metabolic process and mitochondrial respiratory chain complex assembly, whereas miR-7704 downregulated insulin, ERBB, and small GTPase-mediated signaling pathway-related genes. Of particular interest, combining the miR expression changes with their annotated functions, we found a regular pattern, i.e., the miRs upregulated in VU (i.e., miR-34a-5p, miR-34c-5p, miR-424-5p, miR-450- 5p, miR-7704, and miR-516-5p) promoted inflammation but inhibited proliferation; whereas the miRs downregulated in VU (i.e., miR-218-5p and miR-96-5p) were required for cell growth and activation (**Figure 6b**). Therefore, these VU-dysregulated miRs might cooperatively contribute to the stalled wound healing characterized with failed transition from inflammation-to-proliferation(Landen et al., 2016).

**Figure 6.**
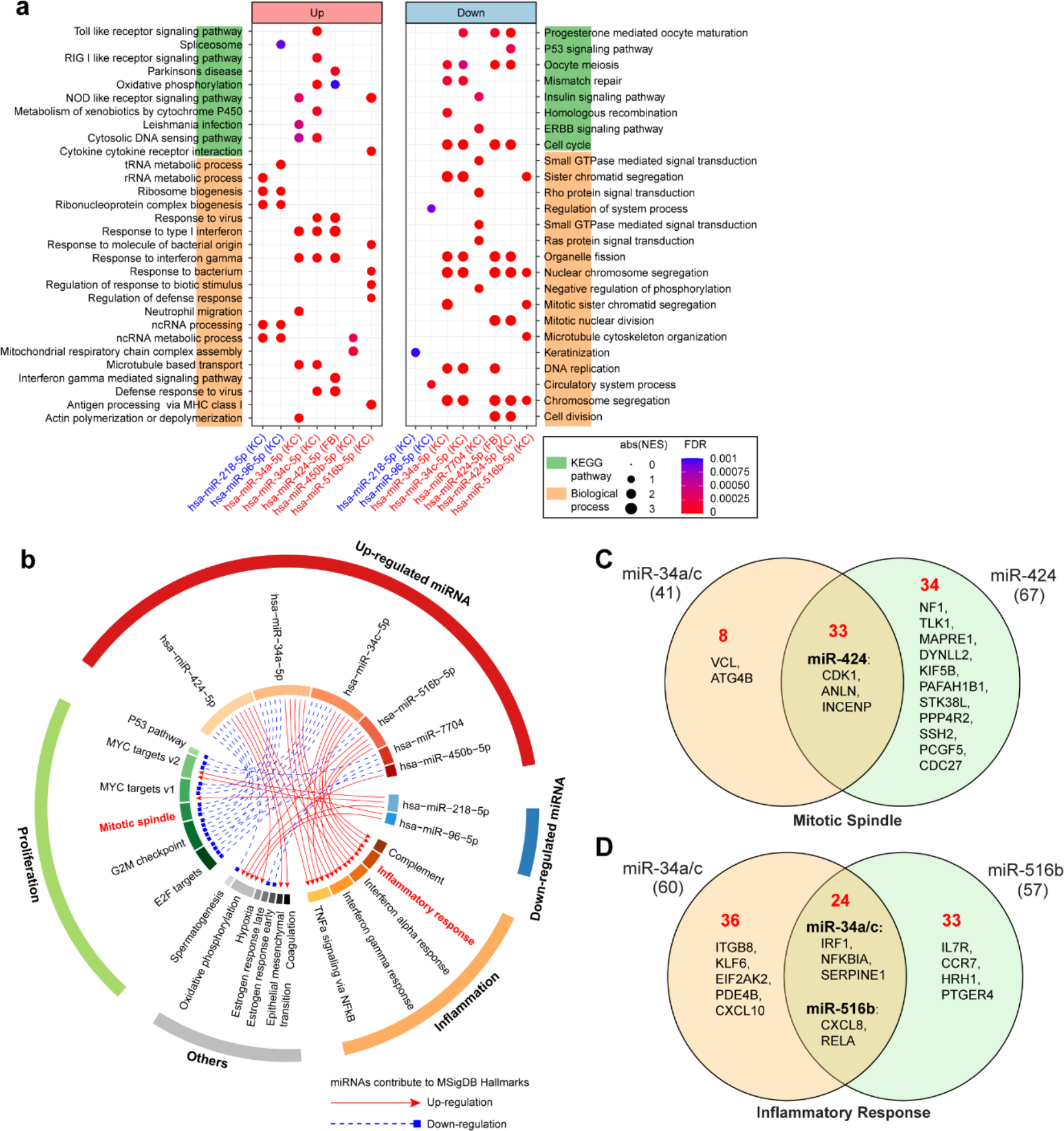
Cooperativity of VU pathology-relevant miRNAs. For the microarray data in keratinocytes (KC) or fibroblasts (FB) with miR-218-5p, miR-96-5p, miR-34a-5p, miR-34c-5p, miR-7704, miR-424-5p, miR-516-5p overexpression, we analyzed KEGG pathways, biological processes **a**, and Molecular Signatures Database (MSigDB) hallmarks **b** enriched by the genes up-or down-regulated by these miRNAs. Venn diagrams show the numbers of the mitotic spindle-related genes regulated by miR- 34a/c-5p and miR-424-5p **c** and the inflammatory response-related genes regulated by miR-34a/c-5p and miR-516b-5p **d** (fold change > 1.2, *P*-value < 0.05) in the microarray analysis of keratinocytes overexpressing these miRNAs. Among the regulated genes, the targets of each miRNA are depicted in the plots.

### Cooperation of miR-34a, miR-424, and miR-516 in regulating keratinocyte proliferation and inflammatory response

To validate miR cooperativity in modulating the key pathological processes in VU, we analyzed proliferation and inflammatory response of keratinocytes overexpressing individual miR or miR combinations. The microarray data gene ontology analysis (**Figure 6b**) showed that three miRs upregulated in VU could suppress the expression of mitotic spindle-related genes (**Figure 6c**): miR- 34a/c-5p reduced the level of 41 mRNAs (including two miR-34a/c targets), while miR-424-5p downregulated the expression of 67 mRNAs (including 14 miR-424 targets). Although 33 mRNAs were commonly regulated by miR-34a/c-5p and miR-424-5p, none of them were co-targeted by these miRs (**Figure 6c**). Similarly, in the cell cycle pathway, miR-34a-5p directly targeted CCND1, CDK6, HDAC1, and E2F3, while miR-424-5p targeted ANAPC13, CCNE1, CDC25B, CDK1, CDKN1B, CHK1, WEE1, and YWHAH, and only CDC23 was co-targeted by both miRs (**Figure 7a**). We thus hypothesized that miR-34a-5p and miR-424-5p might cooperate to impact stronger on cell proliferation by targeting different gene sets within the same signaling pathway. To test this idea, we measured keratinocyte growth by detecting proliferation marker gene Ki67 expression both on mRNA and protein levels. We found that although miR-34a-5p or miR-424-5p alone could reduce Ki67 levels, their combination suppressed stronger Ki67 expression (**Figure 7b–c**, **Figure 7 – supplement 1a** and **additional file 11**). The cooperativity between miR-34a-5p and miR-424-5p in repressing keratinocyte growth was further confirmed by comparing cell growth curves generated with a live cell imaging system (**Figure 7d**, **Figure 7 – supplement 1b** and **Video 1**). Moreover, our microarray analysis showed that the miR-34a- 5p and miR-516b-5p combination extended the list of inflammatory response-related upregulated genes (**Figure 6d**). In line with this, simultaneously overexpressing miR-34a-5p and miR-516b-5p induced a higher inflammatory chemokine CCL20 expression compared to the individual overexpression of each miRNA (**Figure 7e**). In summary, our study identified VU signature miRs, e.g., miR-34a, miR-424, and miR-516, with cooperativity in inflicting more severe pathological changes (**Figure 7f**). These findings open new opportunities of developing wound treatment targeting cooperating miRs with potentially higher therapeutic efficacy and specificity.

**Figure 7.**
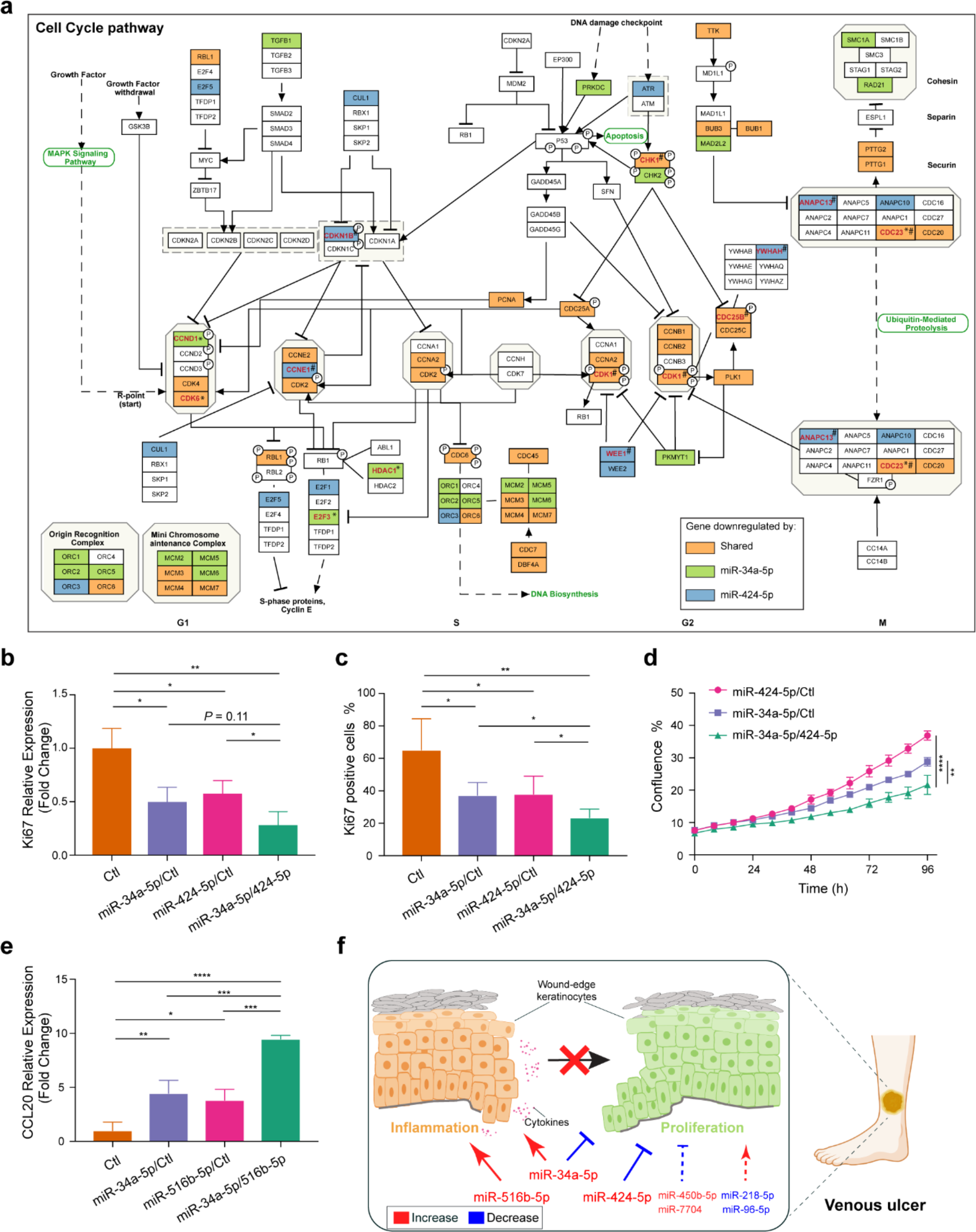
Cooperation of miR-34a, miR-424, and miR-516 in regulating keratinocyte proliferation and inflammatory response. a. In the KEGG cell cycle pathway map, genes regulated by miR-34a-5p or miR-424-5p or both miRNAs as shown in the microarray analysis of keratinocytes are marked with green, blue, and orange colors, respectively. Among the regulated genes, the miRNA targets are highlighted in red and labeled with * for miR-34a targets, ^#^ for miR-424 targets, *^#^ for the gene co- targeted by both miRNAs. Ki67 expression was detected in keratinocytes transfected with miR-34a-5p or miR-424-5p mimics alone or both mimics for 24 hours (n = 3) by qRT-PCR **b** and immunofluorescence staining (n = 5 for Ctl and miR-34a-5p/424-5p, n = 4 for miR-34a-5p/ctl and miR- 424-5p/ctl) **c**. **d** The growth of the transfected keratinocytes (n = 3) was analyzed with a live cell imaging system. **e** qRT-PCR analysis of CCL20 in keratinocytes transfected with miR-34a-5p or miR-516b-5p mimics alone or both mimics for 24 hours (n = 3). **f** Proposed mechanism by which VU-dysregulated miRNAs cooperatively contribute to the stalled wound healing characterized with failed inflammation- proliferation transition. \**P* < 0.05; \*\**P* < 0.01; \*\*\**P* < 0.001 and \*\*\*\**P*<0.0001 by unpaired two-tailed Student’s t-test **b, c,** and **e** and Two-way ANOVA **e**. Data are presented as mean ± SD.

## Discussion

Our genome-wide paired analysis of miR and mRNA expression in human healing and non-healing wounds provides a novel global landscape of the miR regulatory roles in wound biology. A detailed overview of the mRNA expression context at different healing stages or under pathological condition VU allows a more precise understanding about the complex role of miRs in wound repair. The same miR is often described to play different or even opposite roles in different cells, as each cell type has specific gene expression context subjected to the miR-mediated posttranscriptional regulation(Erhard et al., 2014). Thereby, the different mRNA expression profiles in acute or chronic wounds should be considered to understand the precise role of an miR in these distinct contexts. With this aspect in mind, we highlight miRs with their targetome most enriched in the VU mRNA signature, as these miRs display a higher likelihood to regulate pathologically relevant genes. Notably, certain of these miRs did not exhibit the greatest expression change in VU, they would thus be missed with the commonly used strategy that focuses on the top miR expression profiling data changes.

Another strength of our study is the decryption of time-resolved miR-mRNA expression pattern during human skin wound healing, providing a temporal view to our understanding of the functional miR roles. miRs and their target gene expression contexts change dynamically to support different functional needs during wound repair. Defining an miR as “pro-healing” or “anti-healing” requires specifying its temporal expression pattern. For example, continuous expression of a miR that is beneficial for one healing phase but not the other might also lead to deleterious effects.

To understand the molecular mechanisms underlying the miR co-expression patterns, we analyzed the enriched TFs for each miR module with experimentally validated TF-miR regulation data(Tong et al., 2019). This analysis led us to important TFs, such as BMP4(Botchkarev, 2003; Lewis et al., 2014) and GATA3(Kaufman et al., 2003; Kurek et al., 2007) that play fundamental roles in skin development and postnatal remodeling, as well as KLF4, crucial for establishing skin barrier function(Segre et al., 1999). Notably, both GATA3 and KLF4 are reportedly downregulated in human VU(Stojadinovic et al., 2014; Stojadinovic et al., 2008). Our study confirms these findings and provides a novel insight, showing that the loss of these TFs might contribute to VU pathology through their regulated miRs.

Numerous miRs that reportedly regulate wound healing in animal models also surfaced in our study, supporting the robustness of our profiling data and bioinformatics analysis. Thereby, our data would be potentially helpful to evaluate the clinical relevance of these miRs. For example, miR-34a/c reportedly enhance keratinocyte inflammatory response, while suppressing proliferation and migration in cultured cells and mouse wound models(Wu et al., 2020). miR-34a was also identified as one of the most induced miRs in diabetic foot ulcer fibroblasts. Induction of miR-34a together with miR-21-5p and miR-145-5p inhibits fibroblast movement and proliferation, whereas activates cell differentiation and senescence(Liang et al., 2016). In this study, we described that miR-34a/c were specifically upregulated in VU, whereas their levels during wound repair remained relatively low and stable, suggesting their specific role in wound pathology. miR-34 targets were enriched in the M5 mRNA module, containing genes upregulated in the inflammatory phase of wound healing but downregulated in VUs. The VU- relevant miR-34 targetome identified in this study would be potentially useful for determining the precise role of miR-34 in VU pathology. Our current findings in human samples together with previous functional data(Liang et al., 2016; Wu et al., 2020) suggest that miR-34 inhibition along with modulation of additional deregulated miRs might be a promising VU treatment approach.

In addition, certain of these VU-related miRs are involved in skin-related functions but have not yet been linked to wound healing. For example, miR-218-5p regulates hair follicle development(Zhao et al., 2019), inhibits melanogenesis(J. Guo et al., 2014), and enhances fibroblast differentiation(F. Guo et al., 2014). miR-7704 was identified as an exosomal miR produced by melanocytes(Shen et al., 2020). miR-424- 5p suppresses keratinocyte proliferation(Ichihara et al., 2011) and cutaneous angiogenesis(Nakashima et al., 2010; Yang et al., 2017). Moreover, our VU-related miR list (**Figure 4b**) also contains miRs without prior knowledge in their role either in skin or wound healing, e.g., miR-517a-3p, miR-517b-3p, miR-516b-5p, miR-512-3p, and miR-450-5p. It would be highly interesting to examine the role of these miRs in VU. Overall, our dataset can serve as a valuable reference for prioritizing clinically relevant miRs for further functional studies.

Moreover, we studied the relationships between the dysregulated miRs in VU, regarding their target repertoire and biological functions and identified miRs that could act cooperatively. Such knowledge is required for developing combined miR therapeutics with increased specificity and efficacy(Lai et al., 2019). In the miR-target networks underpinning VU (**Figure 4 – figure supplement 1** and **2**), we identified a few putative cooperating miR pairs/clusters that were co-expressed and shared multiple common targets, including the upregulated miR-34a/c together with miR-424-5p and miR-7704, as well as the downregulated miR-218-5p and miR-96-5p in VU. Furthermore, although not sharing targets, the majority of the VU-dysregulated miRs could still regulate the common biological processes coordinately. For example, the miRs upregulated in VU (e.g., miR-34a/c-5p, miR-424-5p, miR-450-5p, miR-7704, and miR-516-5p) promote inflammation but inhibit proliferation; whereas the miRs downregulated in VU (e.g., miR-218-5p and miR-96-5p) are needed for cell growth and activation. As a combined consequence, this VU-miR signature could disrupt the swift transition from inflammation to proliferation (**Figure 7f**). The failure of this phase transition represents a core pathology of chronic wounds(Eming et al., 2014; Landen et al., 2016). Our findings open the possibility of developing innovative wound treatment targeting multiple pathologically relevant cooperating miRs to attain higher therapeutic efficacy and specificity.

Based on the integrative small and long RNA-omics analysis of human wound tissues, we have developed an openly available compendium (https://www.xulandenlab.com/humanwounds-mirna-mrna) for the research community. This novel, rich resource enabled us to gain a network view of miR-mediated gene regulation during human physiological and pathological wound repair in vivo. With the same sequencing datasets, we have also analyzed circular RNA expression and their potential interaction with miRs and miR targets(Toma et al., 2021), which results can be queried at https://www.xulandenlab.com/humanwounds-circrna. These efforts result in many testable hypotheses for future studies elucidating gene expression regulatory mechanisms underpinning tissue repair.

A limitation of our study is the lack of cell type specific miR expression data. Certain gene expression changes detected in the tissue biopsies might be due to the changes of cellular compositions. To rule out this possibility, we validated miR-mediated gene regulation in individual skin cell types (i.e., keratinocytes and fibroblasts) for several miRs surfaced in our analysis. However, to systemically differentiate the gene expression regulation occurring in an individual cell from the altered cellular composition in wound tissues requires future studies using single-cell small RNA sequencing, which technology still remains challenging to be used at a scale for analyzing complex dynamics of tissue, such as human skin and wounds, as it needs extensive cell handing and therefore has only been applied to few cells(Nielsen & Pedersen, 2020). As new technologies for higher cellular resolution miRNA analyses emerge, we hope that such approach will be feasible in a near future.

## Conclusion

In conclusion, this genome-wide, integrative analysis of miR and mRNA expression in human skin and wound tissues reinforce and extend the evidence about the functional role of miRs in wound repair and their therapeutic potential for chronic wound treatment. By combining miR expression patterns with their specific target gene expression context, we identified miRs highly relevant to VU pathology. This rigorous and in-depth molecular characterization of human wound tissues adds a novel dimension to our current knowledge mostly relying on non-human models and would serve as a unique platform and valuable resource for further mechanistic studies of miRs with a high translational potential.

## Data availability

Raw data of small RNA sequencing, long RNA sequencing and microarray performed in this study have been deposited to NCBI’s Gene Expression Omnibus (GEO) database under the accession number GSE174661 (Reviewer access with a token ’cvmpqccylxgnzgj’) and GSE196773, respectively. In addition, the analyzed dataset is presented with an online R Shiny app and can be accessed through a browsable web portal (https://www.xulandenlab.com/humanwounds-mirna-mrna). The analysis source code is available at https://github.com/Zhuang-Bio/miRNAprofiling.

## Funding and Acknowledgements

We express our gratitude to the patients and healthy donors participating in this study. We thank Mona Ståhle, Desiree Wiegleb Edström, Peter Berg, Fredrik Correa, Martin Gumulka, Mahsa Tayefi for clinical sample collection; Helena Griehsel for technical support. We thank the Microarray core facility at Novum, BEA, which is supported by the board of research at KI and the research committee at the Karolinska hospital. The computations/data handling was enabled by resources in the projects sens2020010 and 2021/22-701 provided by the Swedish National Infrastructure for Computing (SNIC) at UPPMAX, partially funded by the Swedish Research Council through grant agreement no. 2018-05973. This work was supported by Swedish Research Council (Vetenskapsradet, 2016-02051 and 2020-01400), Ragnar Söderbergs Foundation (M31/15), Welander and Finsens Foundation (Hudfonden), LEO foundation, Ming Wai Lau Centre for Reparative Medicine, Karolinska Institutet, and R01AR073614 from NIH/NIAMS (to MTC).

## Author Contributions

N.X.L. and P.S. conceived and designed the study. P.S. collected most clinical samples with the assistance of M.A.T.. L.Z., D.L., M.A.T., and X.B. performed the experiments. Z.L. and L.Z. carried out bioinformatics analysis. I.P. and M.T.C contributed to data analyses and interpretation. Z.L., L.Z., and N.X.L. wrote the manuscript, which was commented on by all authors.

---

**Figure 2 - figure supplement 1.**
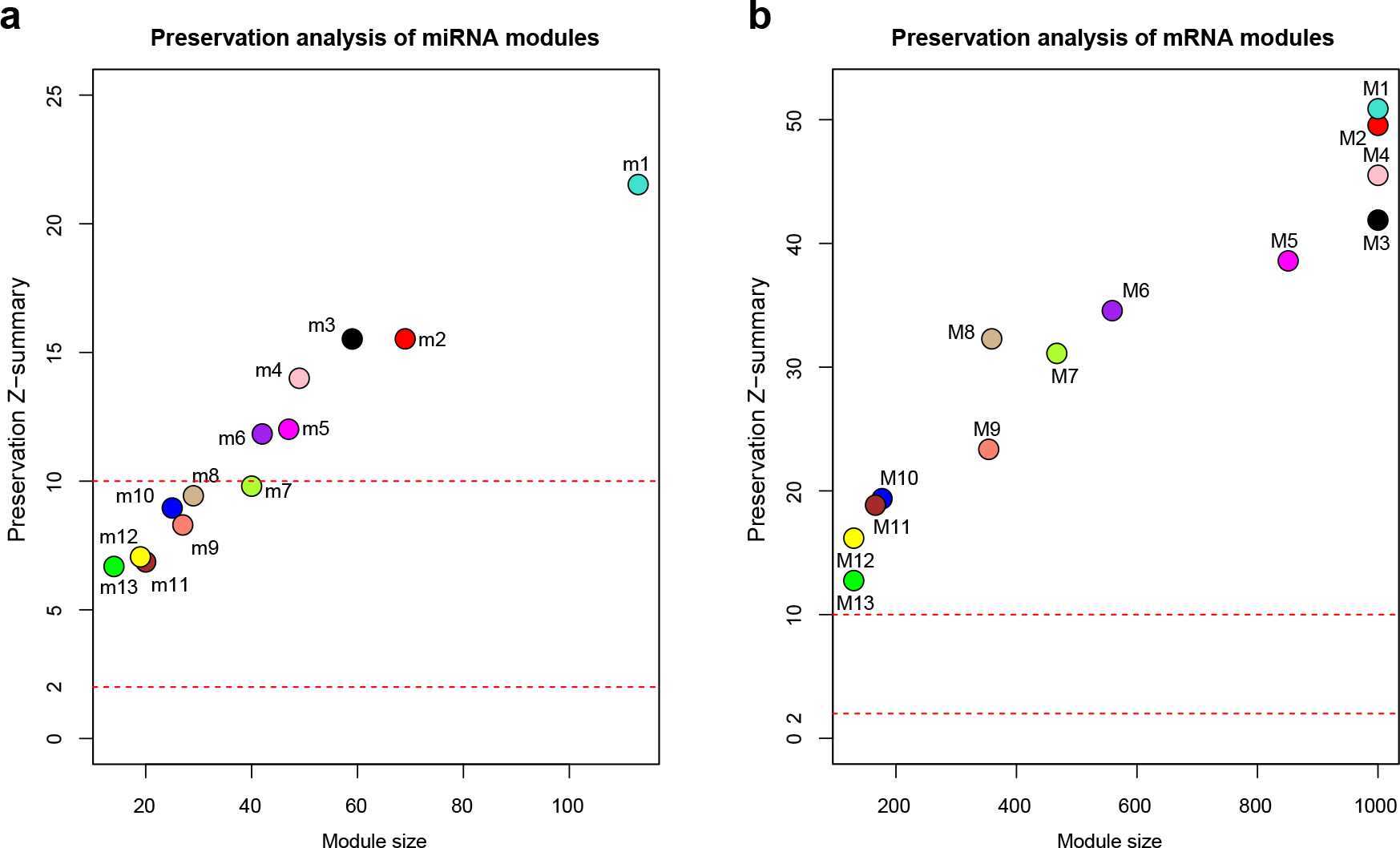
Preservation analysis of miRNA a and mRNA b co-expression modules. The standardized Z-scores were calculated for each module by permutating 200 times using the 20 samples analyzed by RNA-seq as reference and test datasets. Modular preservation is strong if Z-summary > 10, weak to moderate if 2 < Z-summary < 10, no evidence of preservation if Z-summary ≤ 2. miRNA modules.

**Figure 2 - figure supplement 2.**
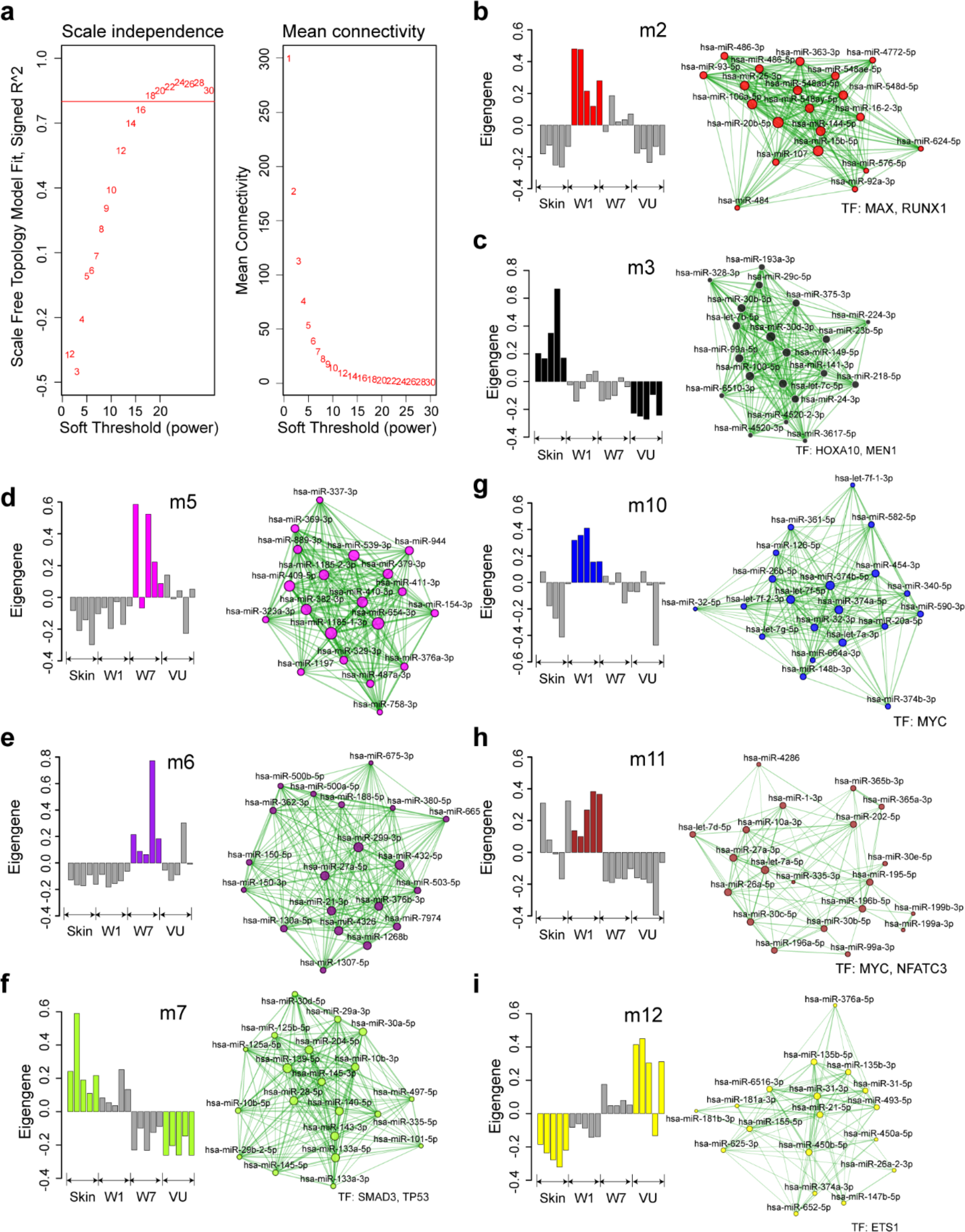
Weighted gene co-expression network analysis (WGCNA) of miRNAs in human skin wound healing. (related to Figure 2a**–d**). **a** The scale-free topology fit index (left) and mean connectivity (right) for various soft-threshold powers. The red line indicates signed R^2^ = 0.8. **b–i** Bar plots (left) depict the ME values across the 20 samples analyzed by RNA-seq and network plots (right) show the top 20 miRNAs with the highest kME values in each module. Node size and edge thickness are proportional to the kME values and the weighted correlations between two connected miRNAs, respectively. Transcription factors (TFs) with their targets enriched in the modules (Fisher’s exact test: FDR < 0.05) are listed below the networks.

**Figure 2 - figure supplement 3.**
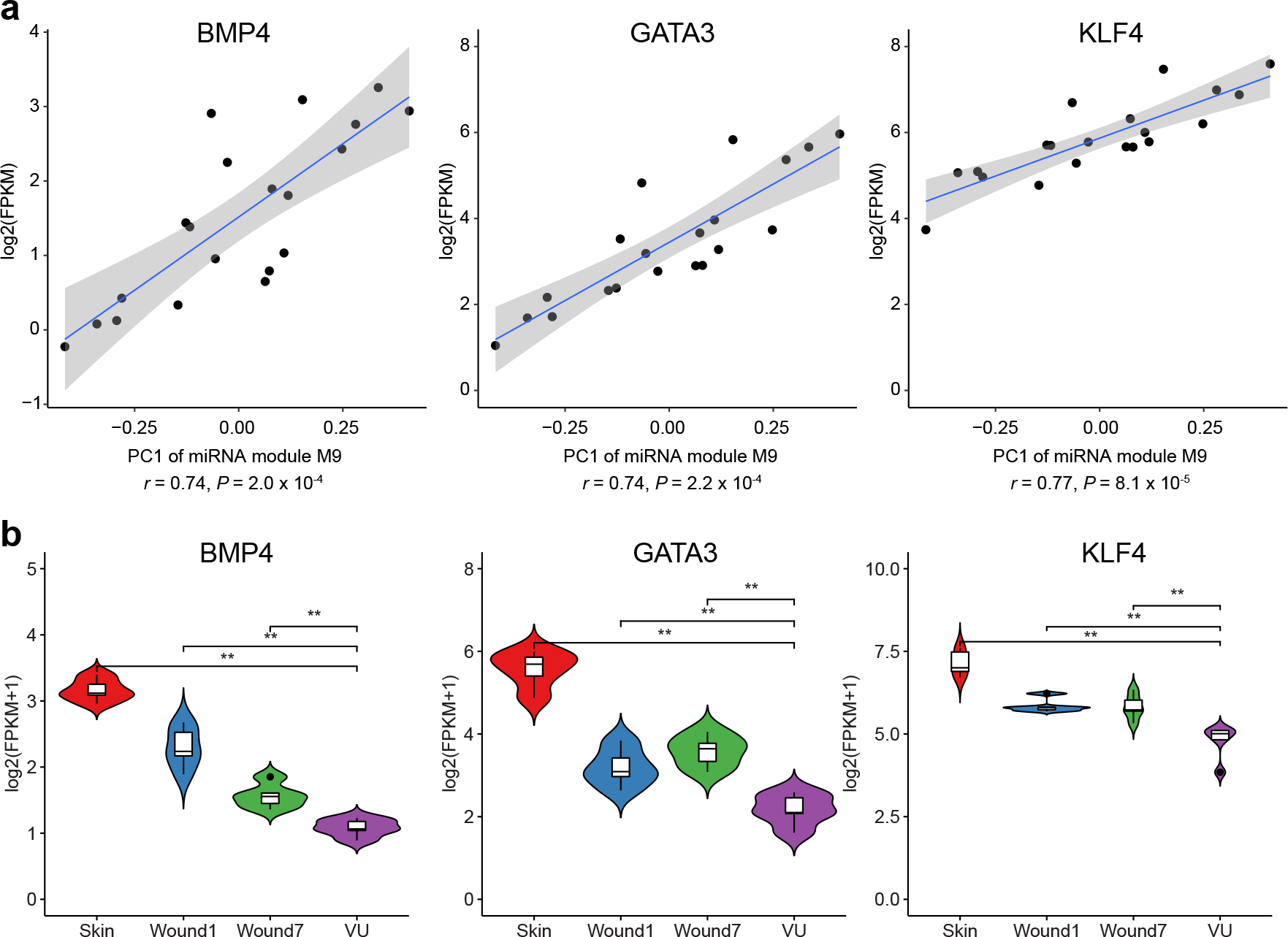
Transcription factors (TFs) with targets enriched in the miRNA m9 module. a. Pearson correlations between the first principal component (PC1) of miRNAs module m9 and the expression of BMP4, GATA3, or KLF4. **b** BMP4, GATA3, and KLF4 expression in the skin, day 1 and day 7 acute wounds from five healthy donors and in five venous ulcers analyzed by RNA-sequencing. Mann-Whitney t-test, \*\**P* < 0.01.

**Figure 2 - figure supplement 4.**
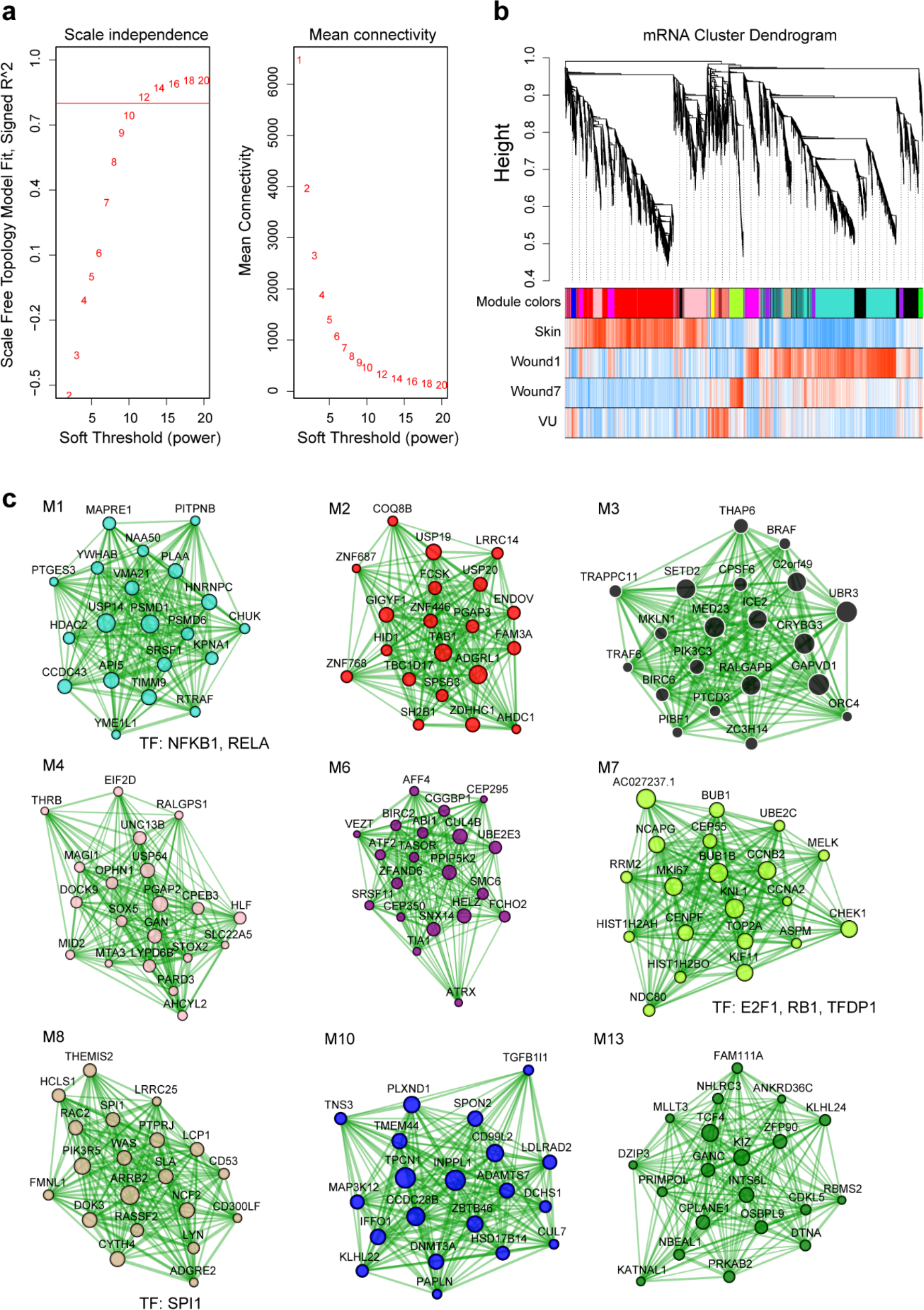
Weighted gene co-expression network analysis (WGCNA) of mRNAs in human skin wound healing. (related to Figure 2e). **a** The scale-free topology fit index (left) and mean connectivity (right) for various soft-threshold powers. The red line indicates signed R^2^ = 0.8. **b** Cluster dendrogram shows mRNA co-expression modules: each branch corresponds to a module, and each leaf indicates a single miRNA. Color bars below show the module assignment (the 1^st^ row) and Pearson correlation coefficients between mRNA expression and the sample groups (the 2^nd^ to the 5^th^ row: red and blue lines represent positive and negative correlations, respectively). **c** Network plots of mRNA modules: the top 20 mRNAs with the highest kME values in each module were plotted in the networks. Node size and edge thickness are proportional to the kME values and the weighted correlations between two connected mRNAs, respectively. Transcription factors (TFs) with their targets enriched in the modules (Fisher’s exact test: FDR < 0.05) are listed below the networks.

**Figure 2 - figure supplement 5.**
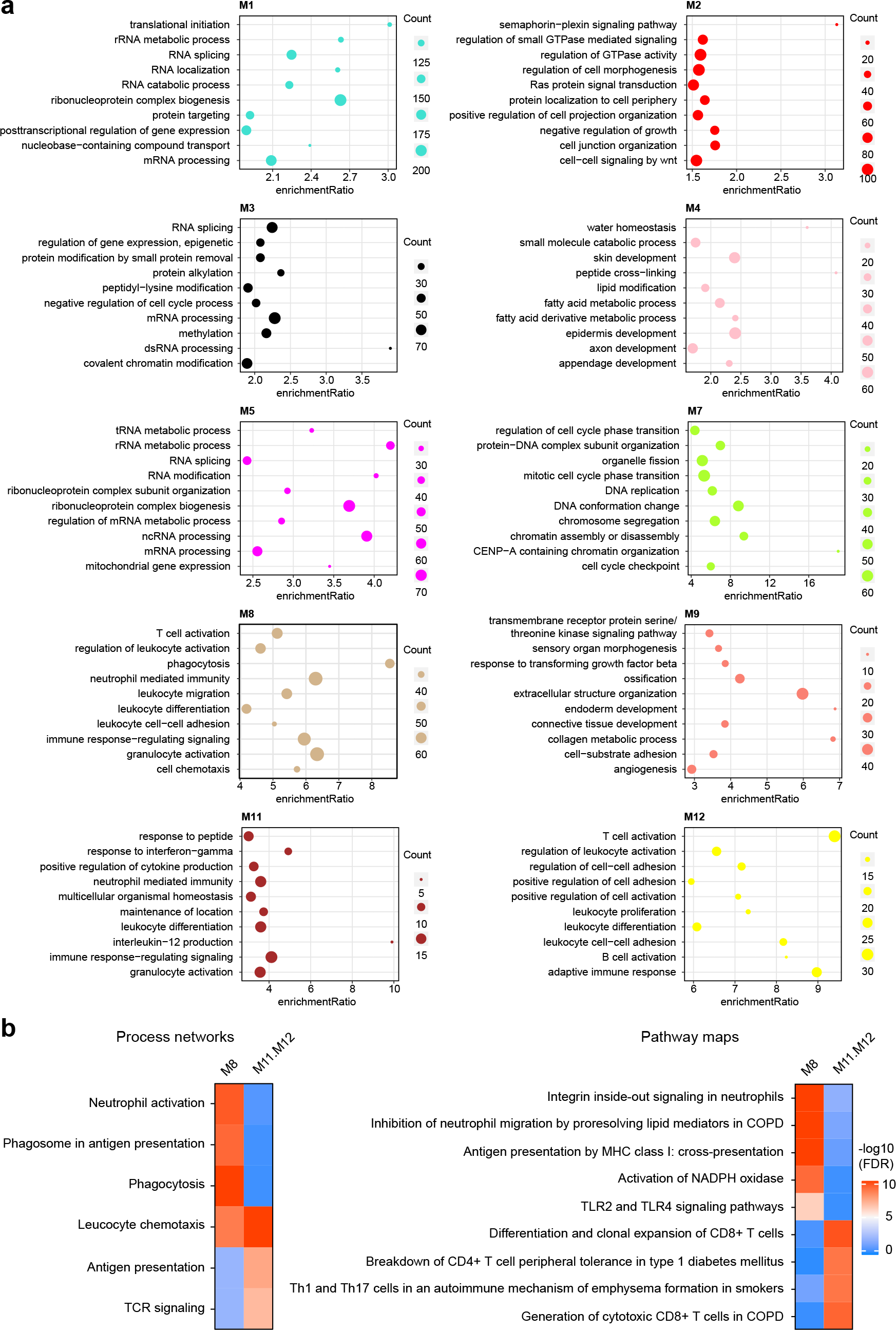
Functional enrichment analysis for mRNA modules. a. The top ten gene ontology (GO) terms with FDR less than 0.05 are shown for each module (related to Figure 2f). **b** MetaCore analysis based on a curated database identified the process networks (left) and pathway maps (right) enriched in the M8 or the combined M11.M12 modules, respectively.

**Figure 4 – figure supplement 1.**
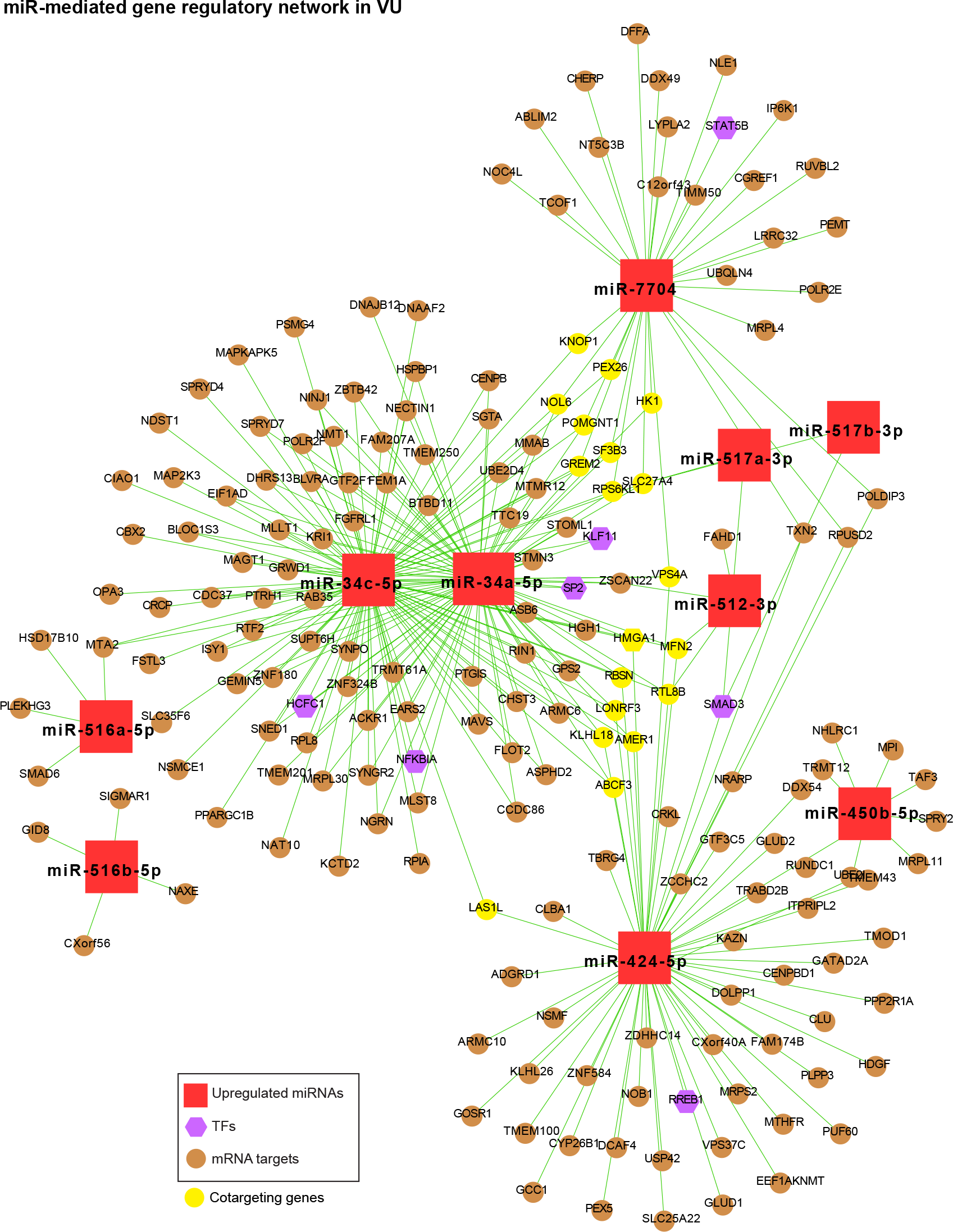
**MiR-mediated gene regulatory network in VU**, which is constructed with the upregulated miRNAs in Figure 4b, the mRNAs predicted as the strongest targets and with anti-correlated expression patterns (Pearson correlation, *P-value* < 0.05 and r < 0) with these miRNAs in human wounds, and the transcription factors (TFs) regulating these miRNAs’ expression from the TransmiR v2 database (related to **Figure. 4b–c** and **Additional file 8**). The mRNAs co-targeted by miRNAs with unrelated seed sequences are highlighted in yellow.

**Figure 4 – figure supplement 2.**
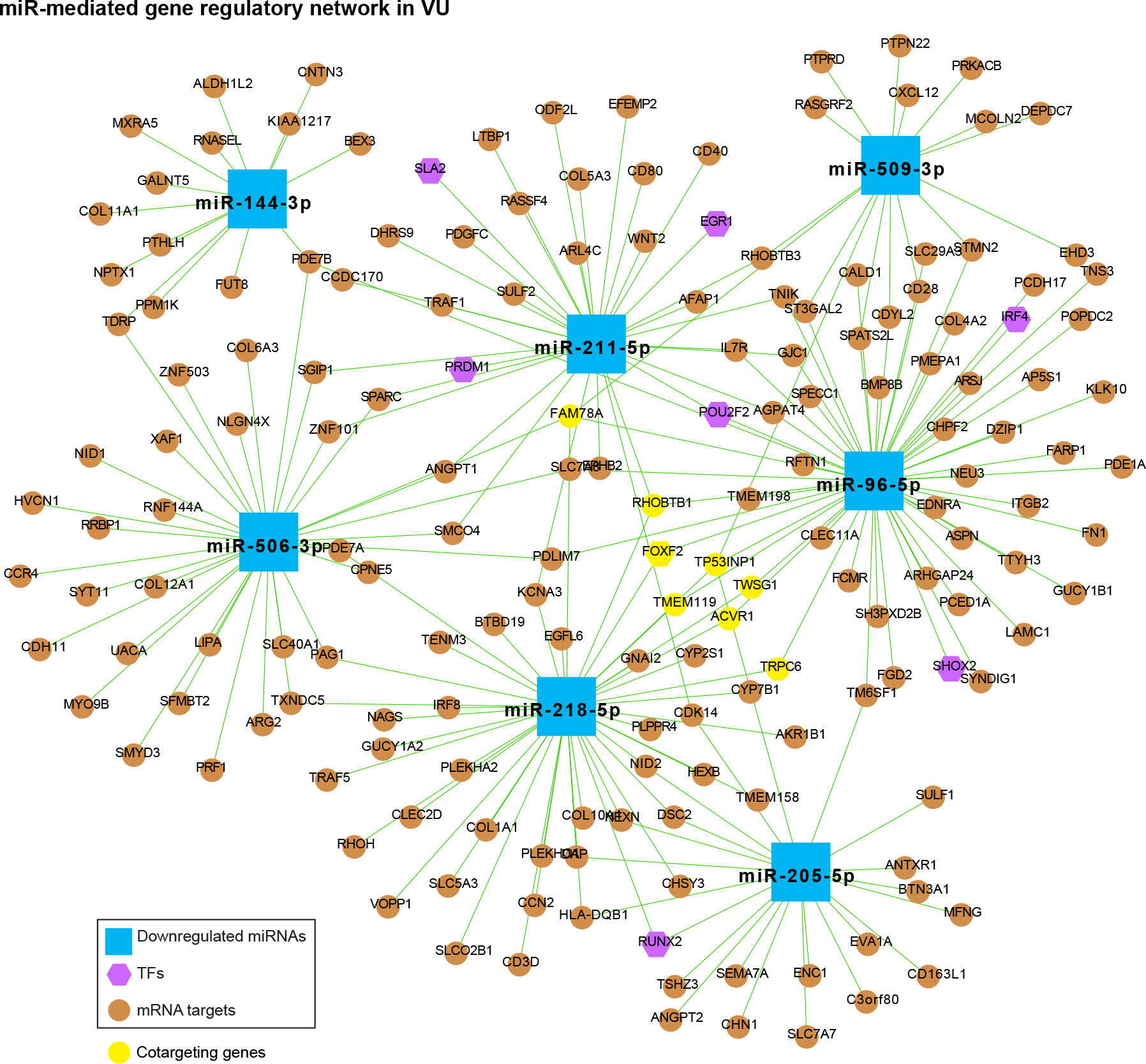
**MiR-mediated gene regulatory network in VU**, which is constructed with the downregulated miRNAs in Figure 4b, the mRNAs predicted as the strongest targets and with anti-correlated expression patterns (Pearson correlation, *P-value* < 0.05 and r < 0) with these miRNAs in human wounds, and the transcription factors (TFs) regulating these miRNAs’ expression from the TransmiR v2 database (related to Figure 4b**–c** and **Additional file 8**). The mRNAs co-targeted by miRNAs with unrelated seed sequences are highlighted in yellow.

**Figure 5 – figure supplement 1.**
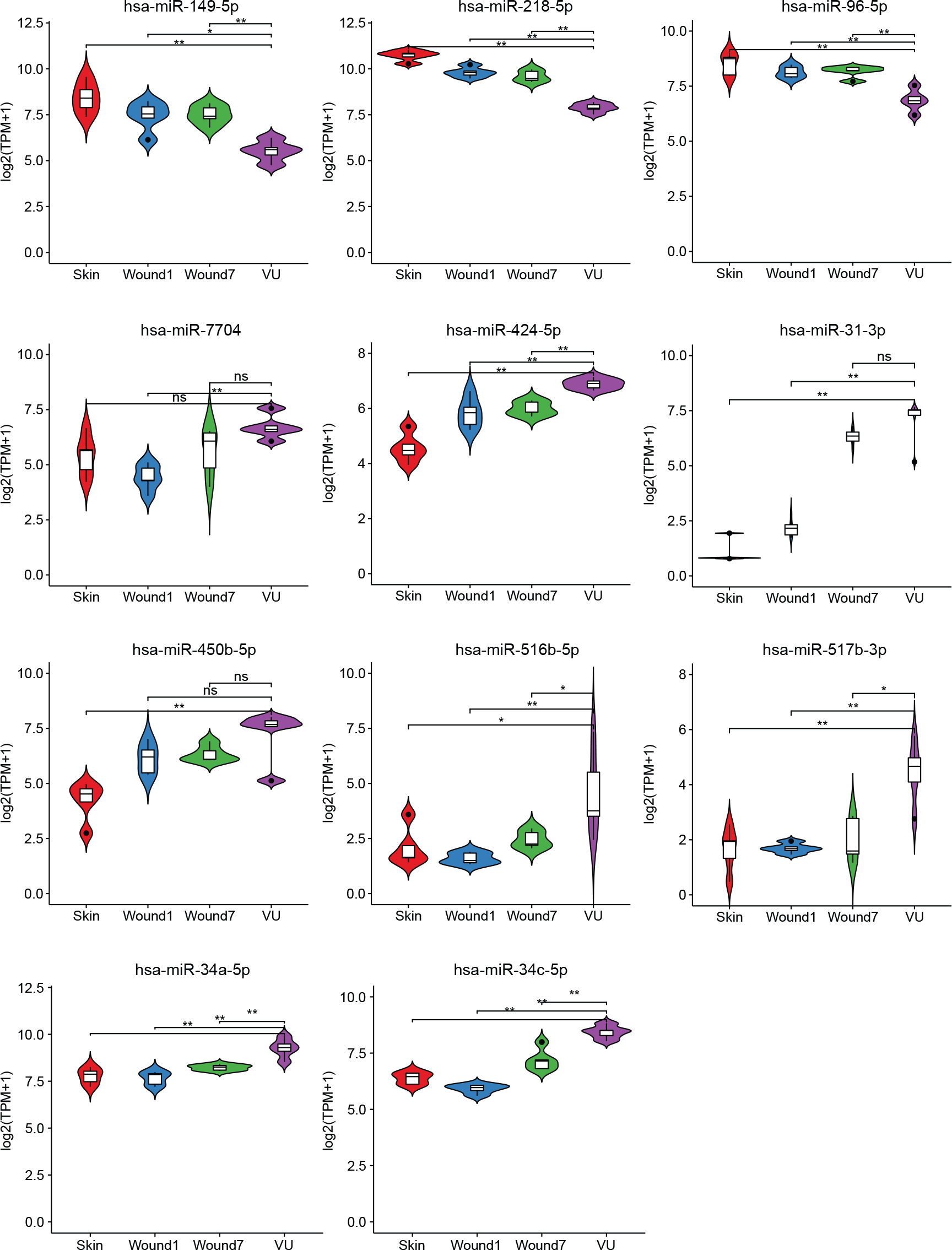
RNA-sequencing results for the miRNAs selected for experimental validation. Mann-Whitney t-test, \**P* < 0.05, \*\**P* < 0.01, ns: not significant.

**Figure 5 – figure supplement 2.**
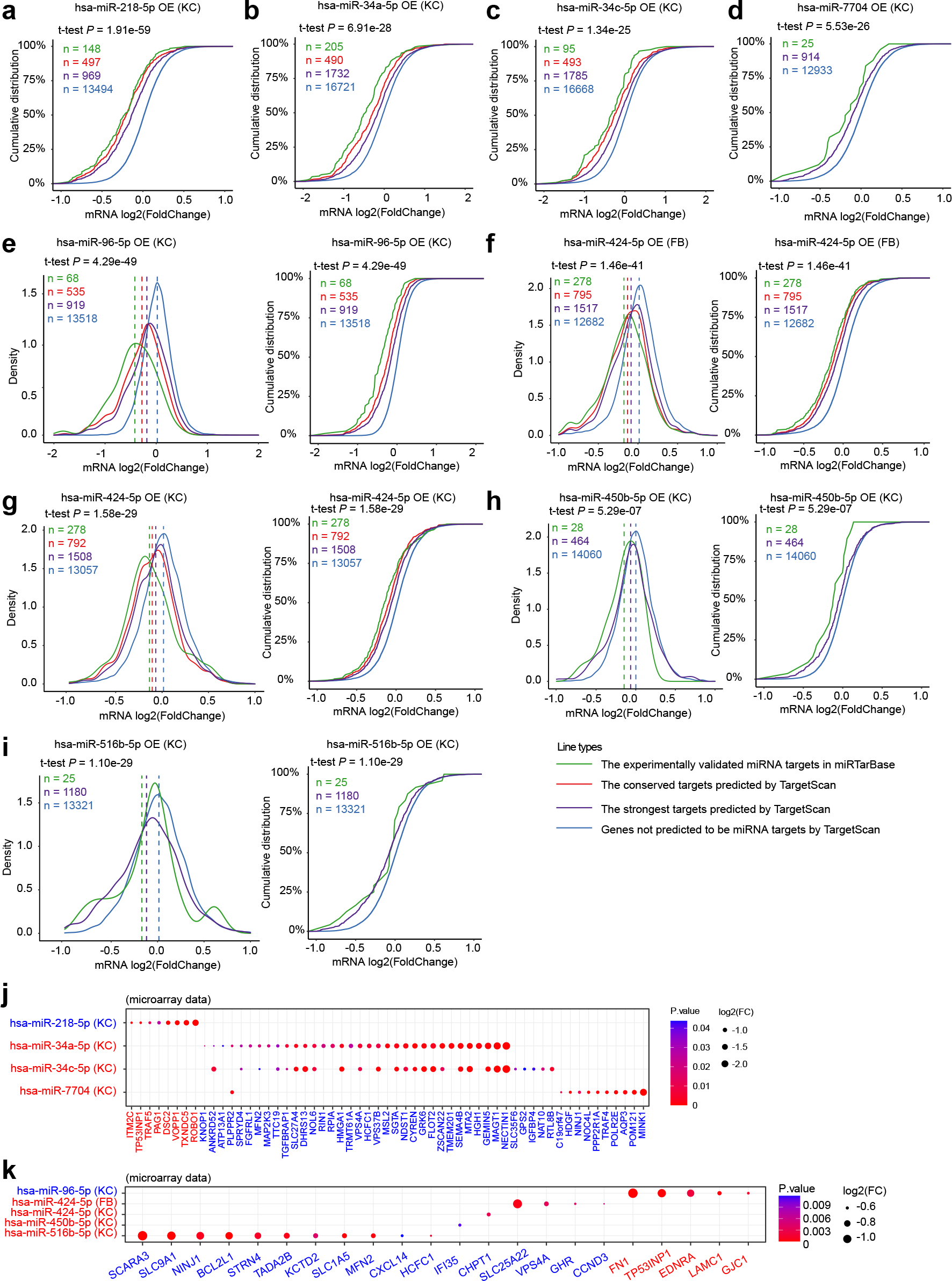
**Experimental validation of miRNAs’ targetome**. Microarray analysis was performed in keratinocytes (KC) or fibroblasts (FB) with miR-218-5p (**a**), miR-34a-5p (**b**), miR-34c- 5p (**c**), miR-7704 (**d**), miR-96-5p (**e**), miR-424-5p (**f, g**), miR-450-5p (**h**), or miR-516-5p (**i**) overexpression (OE). Density plots (left) and cumulative distributions (right in **e–i**) of mRNA log2(fold change) are shown. Wilcoxon t-tests were performed to compare the TargetScan predicted strongest targets (purple) with the non-targets (blue) for each of these miRNAs. The conserved and experimentally validated targets are marked with red and green colors, respectively. Dotted lines depict the average log2(fold change) values for each mRNA group. **j, k** The target mRNAs are significantly changed (fold change < -1.2 and *P*-value < 0.05) by the indicated miRNAs in the microarray analysis.

**Figure 7 – supplement figure 1.**
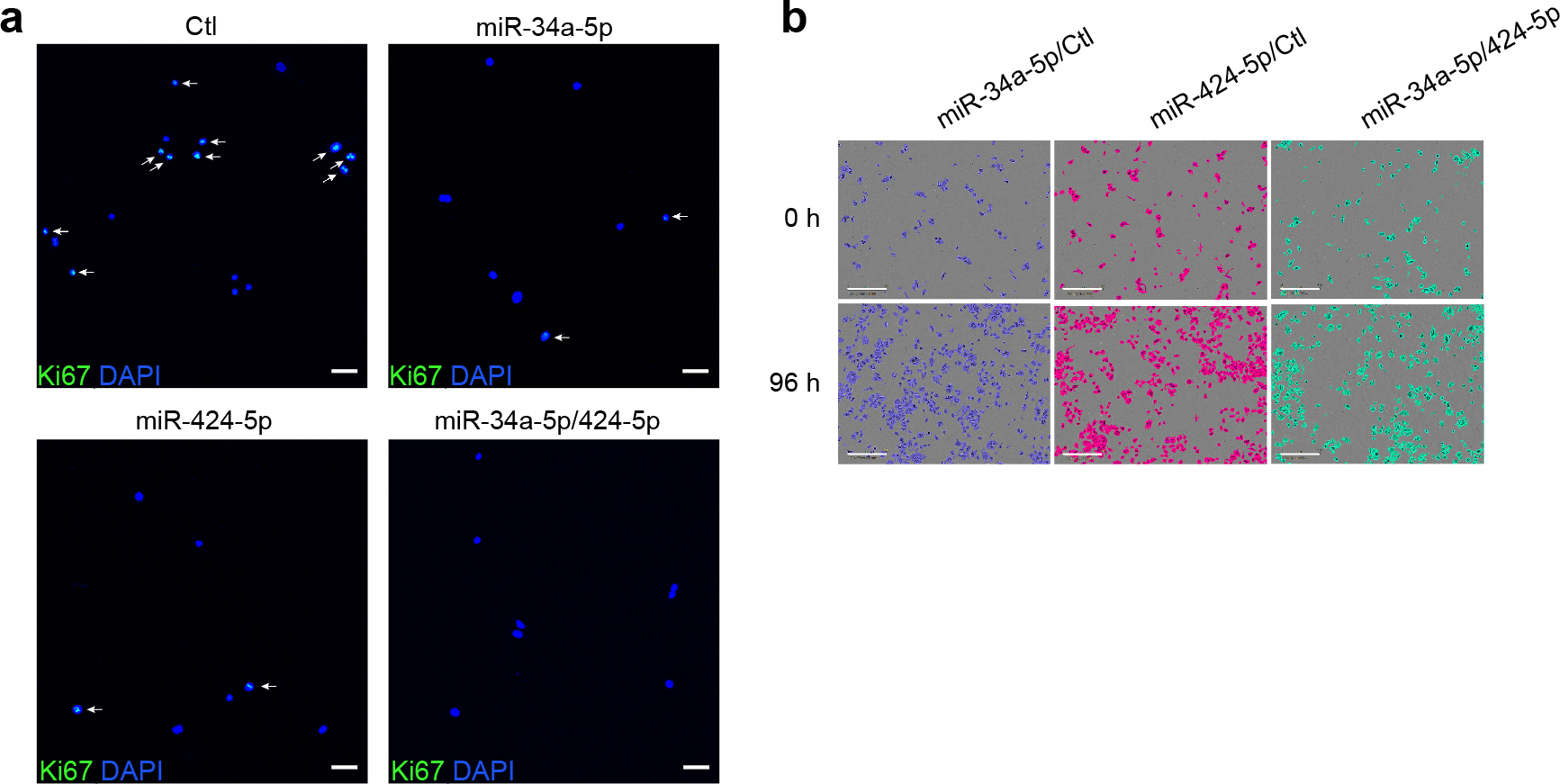
**Cooperation of miR-34a and miR-424 in regulating keratinocyte proliferation**. **a** Ki67 expression was detected in keratinocytes transfected with miR-34a-5p or miR- 424-5p mimics alone or both mimics for 24 hours by immunofluorescence staining. Cell nuclei were costained with DAPI. Ki67+ cell nuclei are highlighted with white arrows in the representative photographs. (Scale bar: 50 μm.) **b** The growth of the transfected keratinocytes (n = 3) was analyzed with a live cell imaging system, and representative photographs of cells at 0 and 96 hours are shown (Scale bar, 300 μm.).

**Video 1**. **Keratinocyte growth was analyzed with a live cell imaging system.** 1, Keratinocytes co-transfected with miR-34a-5p and control mimics. 2, Keratinocytes co-transfected with miR-424-5p and control mimics. 3, Keratinocytes co-transfected with miR-34a-5p and miR-424-5p mimics.

**Additional file 1.** Source data for miRNAs (related to **Figure 1c, d**) and mRNA (related to **Figure 1e**) with expression change specifically in venous ulcers.

**Additional file 2.** Source data for weighted gene co-expression network analysis of miRNAs in wound healing. (related to **Figure 2a, b**)

**Additional file 3.** Source data for the top 20 driver miRNAs of each significant module in WGCNA. (related to **Figure 2c, d** and **Figure supplements 2-2**)

**Additional file 4.** Source data for transcription factors (TF) regulating miRNA expression in each module. (related to **Figure 2c, d** and **Figure supplements 2-2b-2i**)

**Additional file 5.** Source data for weighted gene co-expression network analysis of mRNAs in wound healing. (related to **Figure 2e**)

**Additional file 6**. Source data for transcription factors (TF) with targets enriched in significant mRNA modules. (related to **Figure supplements 2-4c**)

**Additional file 7.** Source data for gene set enrichment analysis for VU-affected DE mRNAs and mRNA modules in the strongest targets of VU-associated DE miRNAs and miRNA modules. (related to **Figure 4a**)

**Additional file 8.** Source data for the individual candidate miRNAs with their targets enriched for the VU mRNA signature. (related to **Figure 4b**)

**Additional file 9.** Source data for experimental validation of miRNAs’ expression in human skin wounds (related to **Figure 5a-i**), enrichment analysis of the experimentally validated miRNA targets for the venous ulcer (VU) gene signature (related to **Figure 5n**), and miRNA targets validated by the microarray and in VU gene signature (related to **Figure supplements 5-2j, 2k**).

**Additional file 10.** Source data for gene ontology analysis of the miRNA-regulated genes in the microarray data. (related to **Figure 6**)

**Additional file 11.** Source data for cooperation of miR-34a, miR-424, and miR-516 in regulating keratinocyte proliferation and inflammatory response. (related to **Figure 7b-e**)

Table S1. Quality control of small RNA sequencing data.

Table S2. Quality control of rRNA-depleted total RNA sequencing data.

