## Supplementary material for "Integrative small and long RNA-omics analysis of human healing and non-healing wounds discovers cooperating microRNAs as therapeutic targets": Table S1 S2

**Table S1. Quality control of small RNA sequencing data.**

| **Groups** | **Sample name** | **Raw**  **reads** | **Q20** | **Q30** | **GC content** | **Clean reads** | **Mapped reads** | **%Mapped rate** |
| --- | --- | --- | --- | --- | --- | --- | --- | --- |
| VU | VU1 | 12352608 | 96.83% | 93.54% | 53.05% | 8592627 | 1478783 | 17.21 |
| VU | VU2 | 10984280 | 96.87% | 93.62% | 52.11% | 7924433 | 1842502 | 23.25 |
| VU | VU3 | 13311104 | 96.85% | 93.61% | 51.73% | 10756575 | 2284593 | 21.24 |
| VU | VU4 | 11251717 | 96.60% | 93.20% | 53.50% | 9426308 | 1847827 | 19.60 |
| VU | VU5 | 16589164 | 96.88% | 93.65% | 51.80% | 14041875 | 2710757 | 19.31 |
| Skin | Skin1 | 17149375 | 95.05% | 89.96% | 51.70% | 15180413 | 5807660 | 38.26 |
| Skin | Skin2 | 11007535 | 95.55% | 91.17% | 50.69% | 9876104 | 6993832 | 70.82 |
| Skin | Skin3 | 10755030 | 95.82% | 91.59% | 51.60% | 9246046 | 4878001 | 52.76 |
| Skin | Skin4 | 10577666 | 96.92% | 93.73% | 52.87% | 9125424 | 3288691 | 36.04 |
| Skin | Skin5 | 14453350 | 97.00% | 93.99% | 51.00% | 12409530 | 6733966 | 54.26 |
| Wound1 | Wound1_1 | 11631465 | 96.76% | 93.38% | 51.62% | 9228914 | 4448460 | 48.20 |
| Wound1 | Wound1_2 | 10202264 | 97.00% | 93.91% | 50.08% | 8894857 | 6258419 | 70.36 |
| Wound1 | Wound1_3 | 10173871 | 96.98% | 93.77% | 50.48% | 8499570 | 5852107 | 68.85 |
| Wound1 | Wound1_4 | 11371432 | 96.90% | 93.71% | 50.77% | 10256444 | 6620450 | 64.55 |
| Wound1 | Wound1_5 | 12469709 | 96.88% | 93.68% | 51.57% | 10400310 | 4957568 | 47.67 |
| Wound7 | Wound7_1 | 11734696 | 96.89% | 93.68% | 51.13% | 9884324 | 4099639 | 41.48 |
| Wound7 | Wound7_2 | 11312106 | 97.08% | 94.07% | 50.42% | 9875669 | 5397518 | 54.66 |
| Wound7 | Wound7_3 | 11326902 | 96.89% | 93.66% | 51.28% | 8587464 | 3942153 | 45.91 |
| Wound7 | Wound7_4 | 11073271 | 96.85% | 93.54% | 52.31% | 8632616 | 2296446 | 26.60 |
| Wound7 | Wound7_5 | 12543302 | 96.85% | 93.58% | 51.70% | 9564730 | 2811963 | 29.40 |

Raw reads, clean reads, and mapped reads are single fragments.

**Table S2. Quality control of rRNA-depleted total RNA sequencing data.**

| **Groups** | **Sample names** | **Total raw**  **reads** | **Total clean reads** | **Clean data rate (%)** | **Input reads** | **Uniquely mapped reads** | **Uniquely mapped**  **rate(%)** |
| --- | --- | --- | --- | --- | --- | --- | --- |
| VU | VU1 | 101622474 | 95087574 | 0.94 | 47543787 | 41697735 | 0.877 |
| VU | VU2 | 101685312 | 92173926 | 0.91 | 46086963 | 35158771 | 0.7629 |
| VU | VU3 | 132592006 | 126190658 | 0.95 | 63095329 | 55771525 | 0.8839 |
| VU | VU4 | 108786868 | 103438880 | 0.95 | 51719440 | 43012289 | 0.8316 |
| VU | VU5 | 108416676 | 101355626 | 0.93 | 50677813 | 41194500 | 0.8129 |
| Skin | Skin1 | 138065658 | 129299094 | 0.94 | 64649547 | 55300913 | 0.8554 |
| Skin | Skin2 | 102996410 | 97056760 | 0.94 | 48528380 | 38950588 | 0.8026 |
| Skin | Skin3 | 123478422 | 116018720 | 0.94 | 58009360 | 47207545 | 0.8138 |
| Skin | Skin4 | 129515874 | 115240764 | 0.89 | 57620382 | 48834112 | 0.8475 |
| Skin | Skin5 | 109239608 | 103341784 | 0.95 | 51670892 | 44434490 | 0.86 |
| Wound1 | Wound1_1 | 105127142 | 100922792 | 0.96 | 50461396 | 44012941 | 0.8722 |
| Wound1 | Wound1_2 | 104542368 | 100678896 | 0.96 | 50339448 | 44040712 | 0.8749 |
| Wound1 | Wound1_3 | 126002796 | 119469842 | 0.95 | 59734921 | 51566913 | 0.8633 |
| Wound1 | Wound1_4 | 103667154 | 98633466 | 0.95 | 49316733 | 41945529 | 0.8505 |
| Wound1 | Wound1_5 | 111446060 | 106610264 | 0.96 | 53305132 | 45076334 | 0.8456 |
| Wound7 | Wound7_1 | 115447748 | 109997276 | 0.95 | 54998638 | 45453970 | 0.8265 |
| Wound7 | Wound7_2 | 104662588 | 100033286 | 0.96 | 50016643 | 42515374 | 0.85 |
| Wound7 | Wound7_3 | 102504766 | 97652674 | 0.95 | 48826337 | 42134005 | 0.8629 |
| Wound7 | Wound7_4 | 108449990 | 103504388 | 0.95 | 51752194 | 43379024 | 0.8382 |
| Wound7 | Wound7_5 | 129466364 | 123563104 | 0.95 | 61781552 | 51511479 | 0.8338 |

Raw reads, clean reads, input reads, and mapped reads are paired fragments.
